# The emerging H3K9me3 chromatin landscape during zebrafish embryogenesis

**DOI:** 10.1101/2024.03.05.582530

**Authors:** Katherine L. Duval, Ashley R. Artis, Mary G. Goll

## Abstract

The structural organization of eukaryotic genomes is contingent upon the fractionation of DNA into transcriptionally permissive euchromatin and repressive heterochromatin. However, we have a limited understanding of how these distinct states are first established during animal embryogenesis. Histone 3 lysine 9 trimethylation (H3K9me3) is critical to heterochromatin formation and bulk establishment of this mark is thought to help drive large-scale remodeling of an initially naive chromatin state during animal embryogenesis. However, a detailed understanding of this process is lacking. Here, we leverage CUT&RUN to define the emerging H3K9me3 landscape of the zebrafish embryo with high sensitivity and temporal resolution. Despite the prevalence of DNA transposons in the zebrafish genome, we found that LTR transposons are preferentially targeted for embryonic H3K9me3 deposition, with different families exhibiting distinct establishment timelines. High signal-to-noise ratios afforded by CUT&RUN revealed new, emerging sites of low-amplitude H3K9me3 that initiated before the major wave of zygotic genome activation (ZGA). Early sites of establishment predominated at specific subsets of transposons and were particularly enriched for transposon sequences with maternal piRNAs and pericentromeric localization. Notably, the number of H3K9me3 enriched sites increased linearly across blastula development, while quantitative comparison revealed a >10-fold genome-wide increase in H3K9me3 signal at established sites over just 30 minutes at the onset of ZGA. Continued maturation of the H3K9me3 landscape was observed beyond the initial wave of bulk establishment.

**Article Summary:** Eukaryotic genomes are organized into transcriptionally permissive euchromatin and repressive heterochromatin. However, we have a limited understanding of how these distinct states are first established during animal embryogenesis. Here, we define the emerging landscape of the heterochromatic histone tail modification H3K9me3 across zebrafish embryogenesis. Our results uncover distinct kinetics of H3K9me3 establishment and growth, define unique characteristics associated with initial sites of H3K9me3, and demonstrate additional maturation of the H3K9me3 landscape beyond the window of bulk H3K9me3 deposition.

## Introduction

The organization of eukaryotic genomes relies on the fractionation of DNA into transcriptionally permissive euchromatin and repressive heterochromatin. Euchromatin, rich in genes, adopts a relatively open chromatin state, while heterochromatin, with fewer genes and an abundance of repeats, is densely packed (Heitz 1928; Vermaak et al. 2003). These distinct regions can be differentiated cytologically through chromosome banding or transmission electron microscopy and molecularly through the detection of histone tail modifications that are crucial for the establishment and maintenance of specific chromatin states (Heitz 1928; Strahl and Allis 2000; Millán-Zambrano et al. 2022).

Histone lysine 9 trimethylation (H3K9me3) is a canonical histone tail modification associated with deeply repressive heterochromatin. H3K9me3 is predominantly found at repetitive sequences, including transposons and satellite repeats. Loss of H3K9me3 leads to chromatin decondensation and can cause chromosome breakage, inappropriate recombination, and increased transposon movement (Allshire and Madhani 2018; Janssen et al. 2018). Additionally, derepression of repetitive elements may activate inflammatory pathways that typically respond to exogenous RNA viruses (Chiappinelli et al. 2015; Roulois et al. 2015; Chernyavskaya et al. 2017; Rajshekar et al. 2018). H3K9me3 is also detected at certain gene regulatory regions in animals where it is implicated in the control of cell identity (McCarthy et al. 2023).

H3K9me3 methylation is catalyzed by conserved methyltransferases. Heterochromatin protein 1 (HP1) binds H3K9me3 and mediates chromatin compaction through oligomerization, phase separation, and the recruitment of additional repressive proteins (Bannister et al. 2001; Lachner et al. 2001; Allshire and Madhani 2018, Zenk et al. 2021). HP1 also promotes the spreading of heterochromatic domains by recruiting H3K9me3 methyltransferases, forming a positive feedback loop to ensure repression (Bannister et al. 2001; Lachner 2001; Allshire and Madhani 2018). While the maintenance of heterochromatic repression is well-studied, targeting of *de novo* H3K9me3 to specific sequences is less well understood. Multiple mechanisms have been implicated in this process in vertebrate animals, including sequence-specific DNA-binding proteins such as KRAB-ZF proteins, small RNAs including piRNAs, and the HUSH complex (Czech et al. 2018; Seczynska et al. 2022; Rosspopoff and Trono 2023). However, most research has explored the role of these pathways in the germline or using cell culture systems.

During early animal embryogenesis, distinct chromatin states are established through a developmentally regulated process. Following fertilization, animal embryos typically exhibit a highly naïve chromatin landscape characterized by a scarcity of histone tail modifications and a deficiency in higher-order chromatin structure (Eckersley-Maslin et al. 2018; Theis and Harrison 2023). In many animal species, the delineation of heterochromatic/euchromatic boundaries and the emergence of cytologically detectable heterochromatin has been shown to roughly follow the onset of the first major wave of transcription from the embryonic genome, a phenomenon commonly referred to as zygotic genome activation (ZGA) (Eckersley-Maslin et al. 2018; Vastenhouw et al. 2019; Theis and Harrison 2023). Dramatic increases in total H3K9me3 are also noted during this period, suggesting that accumulation of H3K9me3 helps to drive the initial formation of condensed heterochromatin following ZGA (Santos et al. 2005; Rudolph et al. 2007; Mutlu et al. 2018; Wang et al. 2018). Altering the kinetics of H3K9me3 establishment compromises animal development and epigenetic reprogramming (Laue et al. 2019; Burton et al. 2020). However, there remains much to learn regarding the mechanisms that target H3K9me3 and the kinetics of its establishment, especially in the context of the developing vertebrate embryo.

Zebrafish offer a powerful model to better understand how heterochromatic regions first emerge during early embryogenesis (Akdogan-Ozdilek et al. 2020). In particular, the large numbers of externally fertilized, synchronously developing embryos produced following zebrafish mating allow for robust analysis of chromatin states from the one-cell stage onward (Kimmel et al. 1995). In zebrafish, the major wave of ZGA initiates after the 10^th^ cell division between 3 and 3.5 hours post fertilization (hpf), and we have previously described the emergence of condensed chromatin ultrastructure and bulk increases H3K9me3 in the window following ZGA (Laue et al. 2019). Unlike mammalian systems, zebrafish do not undergo a major embryonic wave of erasure and reestablishment of the modified base 5-methylcytosine (5mC), and repetitive sequences remain enriched in 5mC throughout this window of development (Potok et al. 2013; Jiang et al. 2013).

Recent advances in chromatin profiling by Cleavage Under Targets and Release Using Nuclease (CUT&RUN) and its adaptation for use in zebrafish allow for quantitative comparison of histone tail modifications across samples using relatively small numbers of early-stage embryos (Skene et al. 2018; Meers et al. 2019; Akdogan-Ozdilek et al. 2022). In particular, the very high signal-to-noise ratios offered by CUT&RUN provide an enhanced opportunity to detect newly emerging sites of H3K9me3 that might be otherwise masked by background. With the clarity and power offered by this approach, we interrogate emerging H3K9me3 profiles across early stages of zebrafish embryogenesis. Our results identify distinct kinetics of H3K9me3 establishment and growth, define characteristics associated with initial sites of H3K9me3, and demonstrate additional maturation of the H3K9me3 landscape beyond the window of bulk H3K9me3 deposition.

## Material & Methods

### Zebrafish

Zebrafish husbandry and care were conducted in full accordance with animal care and use guidelines with approval by the Institutional Animal Care and Use Committees at the University of Georgia. Zebrafish were raised and maintained under standard conditions in compliance with relevant protocols and ethical regulations. Fertilized eggs were obtained from the zebrafish AB strain. Embryos were reared in system water at 28.5°C and staged according to hours post fertilization (hpf) and morphology (Kimmel et al. 1995).

### CUT&RUN

CUT&RUN and library preparation were performed according to Akdogan-Ozdilek et al. 2022 except to facilitate sample processing, time course CUT&RUN samples of ∼30 embryos per replicate were homogenized and rinsed through a 40-μm cell-strainer rather than manually dechorionated. Samples for confirmation experiments were manually dechorionated and normalized to approximately 8,000 cells/replicate. Spike-in E. coli DNA was used for normalization as described in Skene et al. 2018. Time course analysis was performed using an H3K9me3 antibody from Abcam (#8898), and results were confirmed with H3K9me3 antibodies from Active Motif (#39062) and Diagenode (#C15410056).

### Sequencing Data

CUT&RUN libraries were pooled and sequenced on a NextSeq500 instrument (35 bp paired-end for the initial time course, 75 bp paired-end for confirmation experiments) at the Georgia Genomics and Bioinformatics Core (**Table S1**). Sequencing data generated in this study is available through the NCBI GEO database (GSE256288). piRNA sequencing data used in our analysis was previously reported by Kaaij et al 2013 and is available through the NCBI GEO database (GSE41299).

### CUT&RUN Alignment and Normalization

Code used in data analysis and figure generation for this study is available at https://github.com/Goll-Lab/H3K9me3_Kinetics_2024. Short reads (<20 bp) and adaptor sequences were removed using TrimGalore. Trimmed reads were aligned to the current zebrafish genome assembly (GRCz.11 Ensemble release 98) using STAR with the following paramters: --readFilesCommand zcat –outSAMtype BAM SortedByCoordinate –outSAMmultNmax 1 –alignEndsType EndToEnd –alignIntronMax 1 – alignMatesGapMax 2000 (Dobin et al. 2013). Alignments were then filtered using SAMtools for a MAPQ score of 1 (Danecek et al., 2021; Li et al., 2009) and PCR duplicates were removed using Picard MarkDuplicates (Broad Institute, 2019). CUT&RUN data was then normalized for library size and E. coli DNA spike-in using BEDTools “genomecov” (Quinlan and Hall 2010). This outputs a bedgraph file which was converted to a bed file for peak-calling and converted to a bigwig using UCSC “bedGraphToBigWig” for data visualization. Averaged bigwig files were made using deepTools “bigwigCompare” for genome browser visualization (Ramírez et al. 2016). IgG replicates from all time points were averaged together to generate a background file for peak calling and for genome browser visualization.

### Peak Calling

The Hypergeometric Optimization of Motif EnRichment (HOMER) software package was used to identify peaks over IgG using the following parameters: -style histone -minDist 1000 -gsize 1.5e9 -F 4 (Heinz et al. 2010). Peaks from replicates were then analyzed using ChIPr with parameter -m 2 to identify reproducible peaks present in at least two replicates (Newell et al. 2021).

### Peak Annotation

Genic peak annotation was performed using HOMER “annotatePeaks.pl” with the masked reference genome (Heinz et al. 2010). Peaks within 1000 bps of a genic transcriptional start site (TSS) were associated with that gene and gene ontology analysis of those genes was performed using gProfiler (Raudvere et al. 2019). For transposon peak annotation, BEDTools “intersect” with option -F 0.50 was used to identify transposons that were at least 50% covered by an H3K9me3 peak. The updated TE annotation from Chang et al 2022 was used. For comparison of pre-ZGA and post-ZGA enrichment, pre-ZGA peaks were defined as peaks that were present by 3 hpf, while post-ZGA peaks were defined as those present at 4.5 hpf but not already established by 3 hpf. Pre-vs post-ZGA enrichment ratios were calculated by dividing the total number of peaks defined as pre-ZGA by those defined as post-ZGA.

### Mappability Calculation and Threshold

Mappability of zebrafish genome assembly (GRCz.11 Ensemble release 98) was calculated using GenMap with the following parameters: -K 35 -E 3 (Pockrandt et al. 2020). For analysis of transposon families, individual elements that did not have an average mappability score of at least 0.1 were excluded. Additionally, transposon families with fewer than 10 elements meeting this threshold were excluded from analysis.

### Pericentromere Assignment and Analysis

Pericentromeres were assigned using BLAST to find clusters of the satellite repeat BRSATI, which marks zebrafish pericentromeric DNA (Phillips and Reed 2000; Howe et al. 2013). The largest, and often only, cluster of BRSATI sequences was manually selected for each chromosome except for chromosomes 1 and 14 which had no BRSATI repeats present in the current genome assembly. Then, to account for imperfect assembly of the centromeric regions, BEDTools “slop” was used with the options -b 10 -pct to broaden each region 1000% of its length both upstream and downstream (Quinlan and Hall 2010). To generate null control regions, BEDTools “shuffle” with the options -chrom -noOverlapping was used to generate an equal number of size-controlled null regions on the same chromosome. This was repeated so that each pericentromere had 1000 sets of corresponding null regions. Then peak sets of interest were intersected using BEDTools “intersect” with putative pericentromeres and corresponding null regions to determine if sets of peaks were enriched in pericentromeres or not.

### piRNA Alignment and Analysis

Trimmed reads were downloaded from GEO accession #GSE41299 and aligned to the zebrafish genome (GRCz.11 Ensemble release 98) using STAR with the following paramters: --outSAMmultNmax 1 – alignEndsType EndToEnd –alignIntronMax 1 (Dobin et al. 2013). Alignments were then filtered using SAMtools for a MAPQ score of 1 (Danecek et al., 2021; Li et al., 2009) and replicates were merged for further analysis. BEDTools intersect with the options -c -F 0.3 was used to count the number of piRNA reads overlapping with each TE in the genome (Quinlan and Hall 2010). These counts were then used to calculate TPM and only piRNAs with a TPM of greater than or equal to 0.5 were considered in analysis.

### Data Visualization

Heatmaps and profile plots were generated with deepTools (Ramírez et al. 2016). Genome browser images were taken using the Integrative Genomics Viewer version 2.16.2 (Robinson et al. 2011). Karyoplots were generated in R using the “karyoploteR” package (Gel and Serra 2017). Proportional Venn diagrams were made using with R package “euller” (Larsson and Gustafsson 2018). Bar graphs, box plots, and violin plots were generated in R with ggplot2 (Wickham 2009; Wickham et al. 2019).

## Results

### Genome-wide profiling of H3K9me3 across zebrafish embryogenesis

To define the genome-wide kinetics of H3K9me3 establishment across early zebrafish embryogenesis, we used CUT&RUN to assess H3K9me3 in pools of synchronously staged embryos. Embryos were collected in replicates at 30-minute intervals between 2 and 4.5 hours post fertilization (hpf), corresponding to the developmental window surrounding the major wave of ZGA. Confirmation of developmental stages and synchronicity was achieved by verifying the presence of stage-specific morphological features under a bright field microscope (Kimmel et al. 1995). Data analysis included both spike-in and library-depth normalization to enable quantitative comparisons across samples. Concordance among individual enrichment profiles and PCA plots suggested a high degree of correlation between replicates (**Figure S1**).

### H3K9me3 distribution in late blastula stage embryos

The global H3K9me3 landscape of the zebrafish genome has yet to be defined in detail at any stage of development. To remedy this deficiency, and to provide a framework for investigating the developmental trajectory of H3K9me3 establishment, we began our analysis by assessing the genomic distribution of H3K9me3 in the post-ZGA zebrafish genome. Focus was placed on late dome stage embryos (4.5 hpf), as this is a stage at which bulk H3K9me3 is clearly detectable via western blot and condensed heterochromatin is cytologically visible (Laue et al. 2019).

At 4.5 hpf, genome-wide visualization of H3K9me3 revealed regions of substantially increased H3K9me3 at select internal regions on most chromosomes (**Figure 1A**, **S2**). We hypothesized that internal regions of high H3K9me3 signal would correspond to pericentromeric regions, which are marked by H3K9me3 in many species (Allshire and Madhani 2018). The current assembly of the zebrafish genome (GRCz11) is not fully resolved at highly repetitive sequences, and pericentromeric regions are not annotated. To uncover these regions, we identified chromosomal regions enriched in the zebrafish satellite repeat sequence BRSATI, a known cytological marker of zebrafish pericentromeres (Phillips and Reed 2000). Clear regions of high BRSATI density were identified on most chromosomes, and we hypothesize the absence or relatively small BRSATI regions on remaining chromosomes resulted from gaps in the current reference sequence. We noted that across the genome, most regions with a high density of BRSATI repeats also showed overlapping regions of elevated H3K9me3 (**Figure 1A-B**, **S2**). Quantitative comparisons similarly revealed elevated levels of H3K9me3 at the BRSATI sequences themselves when compared to an equal number of random, non-overlapping, size-controlled null regions on each chromosome (**Figure 1B**).

**Figure 1.**
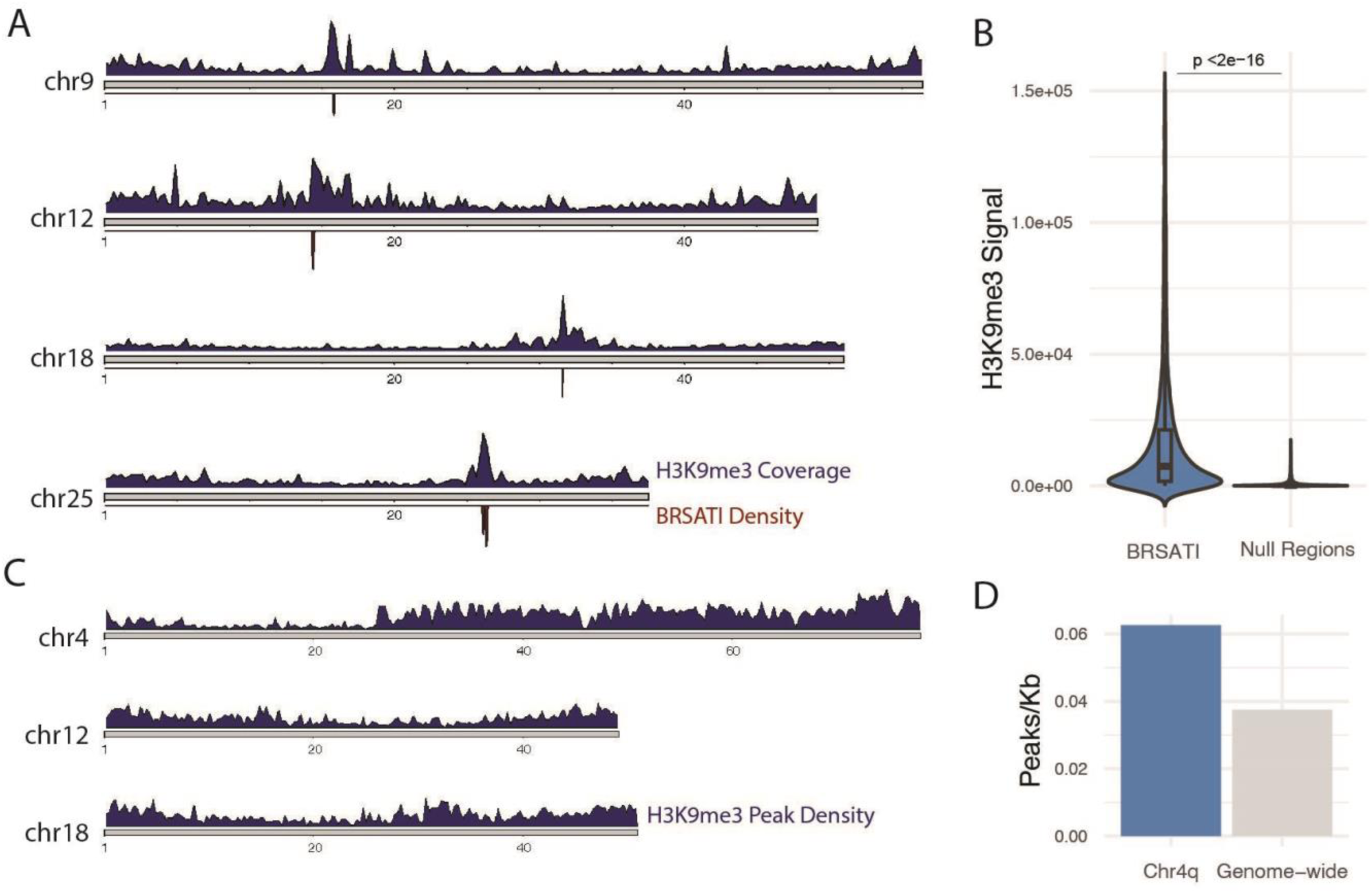
Chromosomal landscape of H3K9me3 in late blastula-stage embryos (4.5 hpf). **A**. H3K9me3 coverage (top, bin size = 250 kb) and pericentromeric repeat BRSATI density (bottom, window size = 50 kb) for representative chromosomes (chr). **B**. Average H3K9me3 signal at BRSATI repeats compared to an equal number of size-controlled regions on each chromosome, values outside 3 standard deviations not plotted for ease of visualization. Two-sample t-test. **C**. H3K9me3 peak density across representative chromosomes (window size 30 kb) **D**. Number of peaks per kilobase (kb) on the long arm of chromosome 4 compared to the number of peaks per kb across the entire genome. For all panels, analysis represents 4.5 hpf data, H3K9me3 signal is averaged across replicates.

We identified ∼50,000 peaks of H3K9me3 enrichment across the genome after excluding BRSATI sequences, where individual peak calling can be challenging. H3K9me3 peak density was similarly distributed on all chromosomes except for the long arm of chromosome 4, which bore a very high density of H3K9me3 peaks across its entire length (**Figure 1C-E**, **S2**). This chromosome arm has been previously hypothesized to be heterochromatinized and is unique in that it exhibits an exceptionally high density of repetitive elements, is late replicating, and has been implicated in zebrafish sex determination in wild strains (Anderson et al. 2012; Howe et al. 2013; Siefert et al. 2017; Chang et al. 2022).

Consistent with a known preference of H3K9me3 for repetitive sequences, we noted a relatively small fraction (∼4%) of H3K9me3 peaks within 1 kilobase (kb) of the transcriptional start site (TSS) of an annotated gene. Among peaks near gene TSSs, roughly two thirds overlapped with transposable elements, with ∼9% being LTR transposons (**Figure S3**). Gene ontology analysis did not reveal any notable categories of genes with H3K9me3 enrichment.

### LTR transposons are preferential targets of H3K9me3 in late blastula-stage embryos

H3K9me3 enrichment at transposable elements is widely observed across diverse eukaryotes (Allshire and Madhani 2018). However, the transposon ecosystem in different organisms can vary significantly. For example, the repetitive portion of mammalian genomes is dominated by LINE and SINE elements, followed by LTR retrotransposons, with few DNA transposons (Pace and Feschotte 2007; Chalopin et al. 2015). In contrast, DNA transposons make up nearly 46% of the zebrafish genome, while LTRs, LINES, and SINEs represent roughly 6%, 4%, and 3% of the zebrafish genome respectively (Chang et al. 2022).

Among the ∼50,000 sites of identified H3K9me3 enrichment in 4.5 hpf embryos, ∼88% were at annotated transposons. Most transposon sequences marked by H3K9me3 were DNA transposons (78%), which is consistent with the heavy bias toward these elements in the zebrafish genome (Figure 2A). Adjusting for abundance, the fraction of DNA transposons enriched for H3K9me3 was quite low, with only 4% of DNA elements associated with H3K9me3 peaks (Figure 2B). At the family level, among roughly ∼1000 annotated families of DNA transposons, we found only 8 (0.79%) where at least 20% of elements were enriched for H3K9me3 (Figure 2C). Among these, we noted all three zebrafish Crypton-H transposon families and 2/76 zebrafish PIF-Harbinger families.

**Figure 2.**
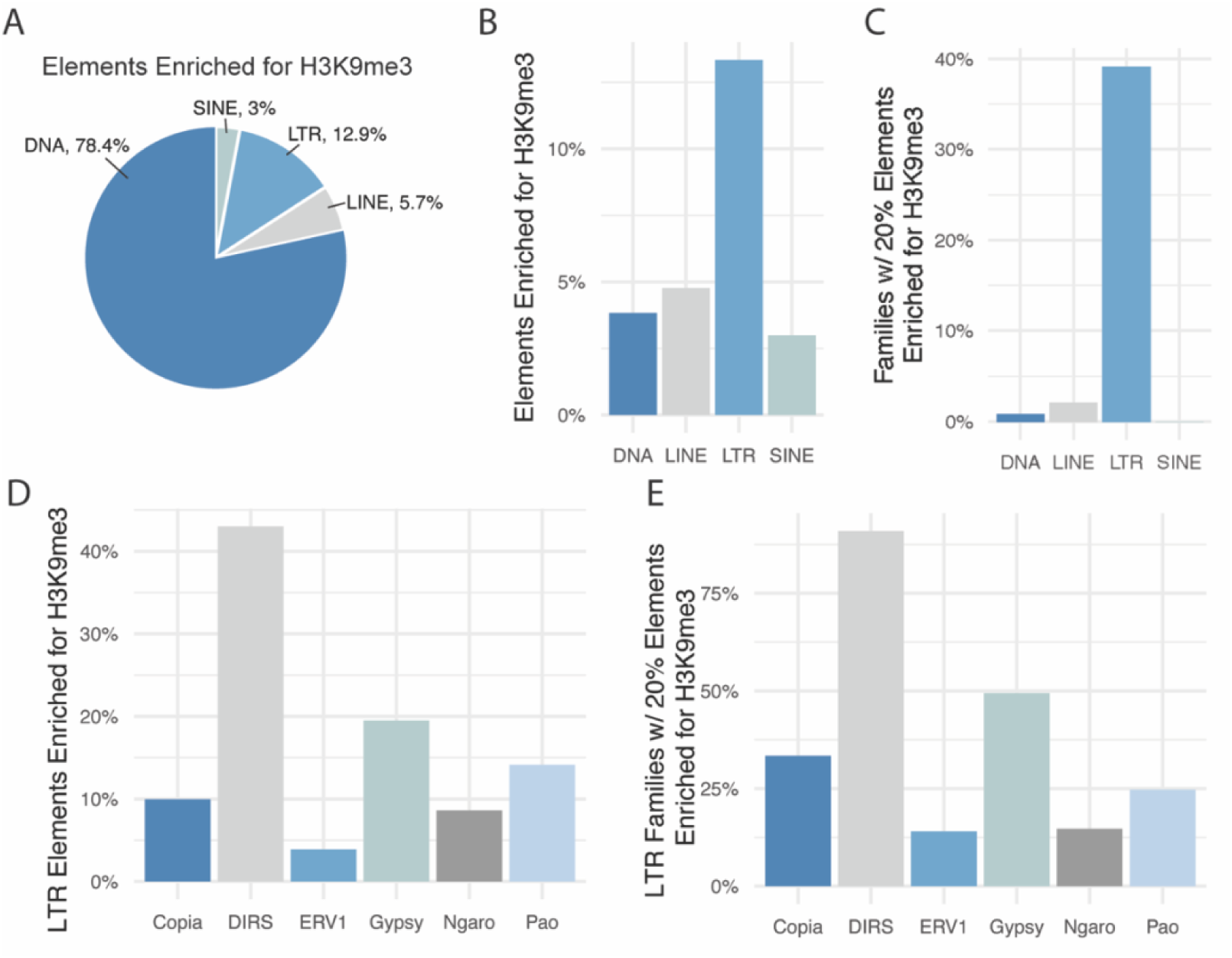
Differential enrichment of H3K9me3 at transposable elements in late blastula-stage embryos **A**. Fraction of total transposable elements enriched for H3K9me3 for each transposon category. **B**. Percent transposable elements enriched for H3K9me3 normalized for abundance. **C**. Percent of transposable element families with at least 20% of individual elements enriched for H3K9me3. **D**. Percent of LTR transposable elements enriched for H3K9me3 relative to abundance. **E**. Percent of LTR families with at least 20% of elements enriched for H3K9me3. For all panels, data reflects 4.5 hpf. Only elements with a mappability score of at least 0.1, and families with at least 10 elements meeting this threshold are considered.

We detected a substantially higher prevalence of H3K9me3 at LTR transposons relative to their genomic abundance, with 13% of LTR transposons enriched in H3K9me3 (Figure 2B). This included 201/514 (39%) LTR transposon families where at least 20% of copies were marked by H3K9me3 (Figure 2C). Among LTR transposons, the most extensive H3K9me3 enrichment was observed for Gypsy (19% of elements, 131/265 families) and DIRS retrotransposons (43% of elements, 10/11 families) (Figure 2D**-E**). We did not observe any families of SINE elements with notable H3K9me3 enrichment, and among LINE elements, only 7/337 (2%) families showed H3K9me3 enrichment at 20% of elements or more (Figure 2C). For DNA, LINE, and LTR transposons, we noted that families that were more extensively enriched in H3K9me3 tended to have shorter median phylogenetic branch lengths, suggesting a bias toward younger elements (**Figure S4**). These younger elements are more likely to have been recently active in the zebrafish genome, potentially making them preferred targets for chromatin mediated silencing.

### Low amplitude H3K9me3 establishment events precede ZGA

To investigate how the H3K9me3 landscape develops over early embryogenesis, we compared the global H3K9me3 enrichment profile of 4.5 hpf embryos to earlier stages in the CUT&RUN developmental time course. Consistent with western blot data examining bulk H3K9me3 (Laue et al. 2019), initial analysis suggested limited H3K9me3 signal in the embryonic genome before the major wave of ZGA, which initiates between 3 and 3.5 hpf (Figure 3A, **C**). However, when pre-ZGA time points were scaled separately, H3K9me3 enrichment became apparent at a subset of sites across the genome (Figure 3B**, D**).

**Figure 3.**
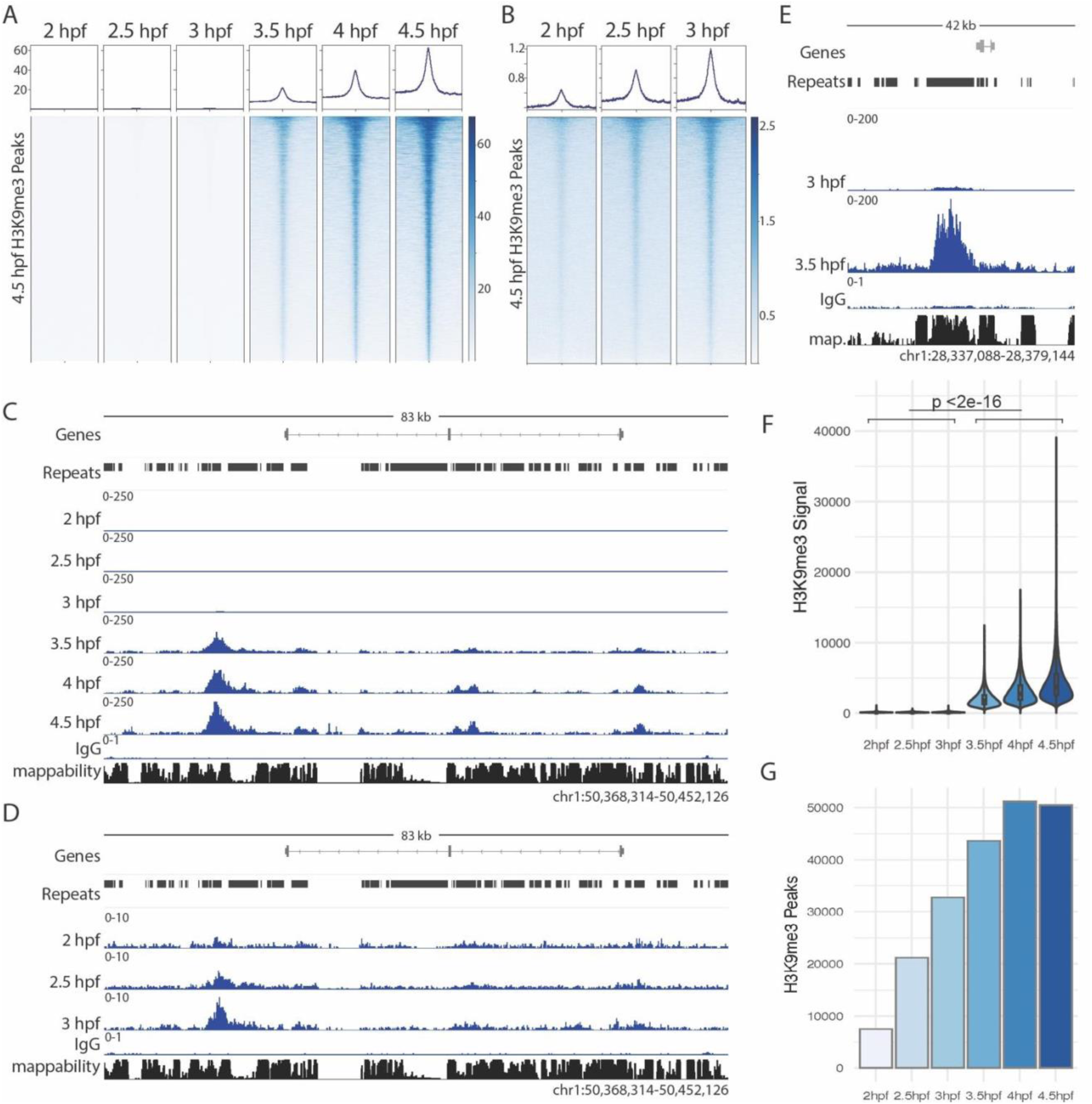
A time course of H3K9me3 enrichment reveals pre-ZGA H3K9me3 establishment **A**. Heatmap of H3K9me3 signal at each timepoint ranked on 4.5 hpf peaks. **B**. Heatmap of H3K9me3 signal from pre-ZGA timepoints in (A) rescaled for increased sensitivity. **C**. Genome browser image of H3K9me3 enrichment with all time points equally scaled. **D**. Genome browser image of pre-ZGA time points for the same region shown in (C) rescaled to enable visualization. **E**. Genome browser image demonstrating the rapid increase in H3K9me3 signal between 3 hpf and 3.5 hpf with tracks scaled equally **F**. Average H3K9me3 signal in called peaks at each time point, values outside 5 standard deviations not plotted for ease of visualization. Two-sample t-test. **G**. Total number of peaks called at each time point. For all panels, data represents H3K9me3 signal averaged across time point replicates.

To rule out the possibility that these putative pre-ZGA enrichment sites resulted from contamination due to somatic cells adhering to the embryo chorion during sample preparation, we repeated H3K9me3 CUT&RUN using cell-count normalized samples derived from manually dechorionated 2.5 hpf embryos.

This analysis again revealed the presence of low amplitude H3K9me3 enrichment sites at 2.5 hpf, with sites of enrichment mirroring those observed in the original time course (**Figure S5**). We were also able to detect these pre-ZGA H3K9me3 sites of enrichment using two additional commercially available H3K9me3 antibodies, further supporting their authenticity (**Figure S5**).

<u>H3K9me3 establishment at select sequences precedes bulk accumulation in the early embryo</u> Consistent with the separate rescaling required to visualize pre-ZGA H3K9me3 peaks, quantitative comparisons between 3 and 3.5 hpf time points confirmed a >10 fold increase in the average amplitude of H3K9me3 peaks specifically during this 30-minute window, with large amplitude increases observed across the genome (Figure 3E-F**, S6**). This temporally restricted surge in amplitude contrasted with an observed linear growth in the total number of detected H3K9me3 peaks during the period between 2 and 4 hpf (Figure 3G). These vastly different temporal trajectories demonstrate that H3K9me3 establishment and signal growth are not fully coincident in the early zebrafish embryo.

We found that once established, the majority of detected H3K9me3 enrichment sites persisted through the end of the time course, with 84% of enrichment sites detected at 2.5 hpf still present at 4.5 hpf (**Figure S6**). Within the time course, we did observe some H3K9me3 enrichment sites that were only present at one time point or that appeared and disappeared within our timeline (**Figure S6**). These events were much rarer and could be a result of technical noise. However, given the presence of these enrichment sites in multiple replicates, it is also possible that they represent sites of short-lived H3K9me3 accumulation that are lost during developmental progression.

### Pre-ZGA H3K9me3 is biased toward LTR transposons

We noted that pre-ZGA peaks accounted for roughly one third of H3K9me3 peaks present in the 4.5 hpf zebrafish embryo (**Figure S6**). Comparing these low amplitude peaks detected before ZGA (pre-ZGA peaks) to peaks that were first detected after ZGA (post-ZGA peaks) revealed notable biases, suggesting that unique targeting mechanisms predominate in these two developmental windows. At LTR transposons, H3K9me3 enrichment was significantly biased toward pre-ZGA establishment, while DNA transposons were more likely to gain H3K9me3 after ZGA (Figure 4A**, S7**). Among LTR transposons, there was a significant bias towards pre-ZGA H3K9me3 at conventional Gypsy and Pao retroelements. Similar bias was not observed for tyrosine recombinase encoding DIRS and Ngaro retrotransposons, Copia elements, or ERVs (Figure 4B**, S7**). Additionally, we noted LTR elements with shorter median phylogenetic branch length were more likely to exhibit pre-ZGA H3K9me3 enrichment, suggesting a bias towards early targeting of these younger elements (**Figure S7**).

**Figure 4.**
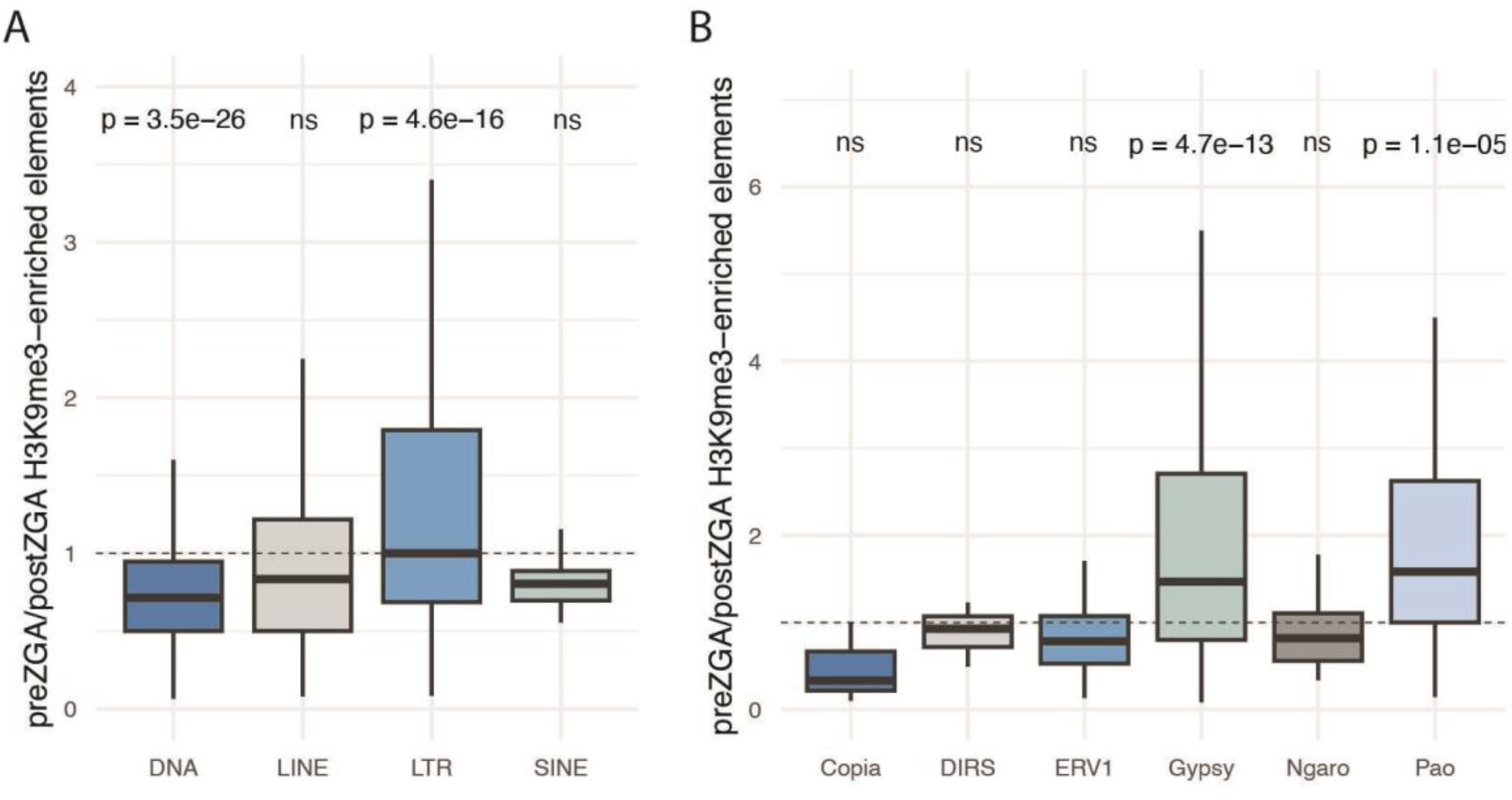
pre-ZGA H3K9me3 establishment at transposons is biased towards LTRs **A**. Box and whisker plot displaying the ratio of H3K9me3-enriched pre-ZGA elements to H3K9me3-enriched post-ZGA elements per transposon family for each category. One sample t-test: mu = 1, indicated by dashed line. **B**. Identical analysis exclusively examining LTR transposons. For all box and whisker plots: center line represents the median, hinges represent 25^th^ and 75^th^ percentiles, and whiskers extend an additional 1.5 times their respective interquartile range. Outliers not plotted. For all panels, only elements with a mappability score of at least 0.1, and families with at least 10 elements meeting this threshold are considered.

In many species, piRNAs are associated with transposon constraint in the germline and maternally derived piRNAs can be transmitted to progeny (Houwing et al. 2007; Houwing et al. 2008; Czech et al. 2018; Guo et al. 2021). In Drosophila, depletion of embryonic piRNAs can lead to heterochromatic derepression and enrichment of embryonic piRNAs has been observed in genomic regions near some sites of early H3K9me3 establishment (Gu and Elgin 2013; Fabry et al. 2021; Wei et al. 2021). Recent preprinted work has also reported maternally loaded piRNAs corresponding to some LTR transposons that are silenced by H3K9me3 in the post-ZGA zebrafish embryo (Guo et al. 2021). Together, these findings implicate piRNAs in the regulation of embryonic heterochromatin. However, their specific targets and relative importance for H3K9me3 establishment versus signal reinforcement remain unclear.

Taking advantage of published piRNA sequencing data (Kaaij et al. 2013), we found a significant enrichment of maternal zebrafish piRNAs corresponding to sites of pre-ZGA H3K9me3 enrichment when compared to sites that were only marked by H3K9me3 post-ZGA (Figure 5A). Further analysis revealed this association was almost exclusively driven by reverse strand piRNA sequences and was strongest for younger LTR elements (Figure 5B, **S7**). Consistent with this analysis, we find that among transposons with pre-ZGA H3K9me3 enrichment, LTR families were most likely to have corresponding maternal piRNAs (Figure 5C**, S7**). This trend is driven primarily by Gypsy and Pao elements, with piRNAs corresponding to the majority pre-ZGA H3K9me3 peaks within each of these families (Figure 5D**, S7**). Together, these findings suggest a role for piRNAs in shaping the pre-ZGA H3K9me3 landscape. At the same time, we also noted many sequences in the genome with corresponding maternal piRNAs that lacked H3K9me3 enrichment, suggesting that piRNAs alone are generally not sufficient to direct pre-ZGA H3K9me3 (Figure 5E).

**Figure 5.**
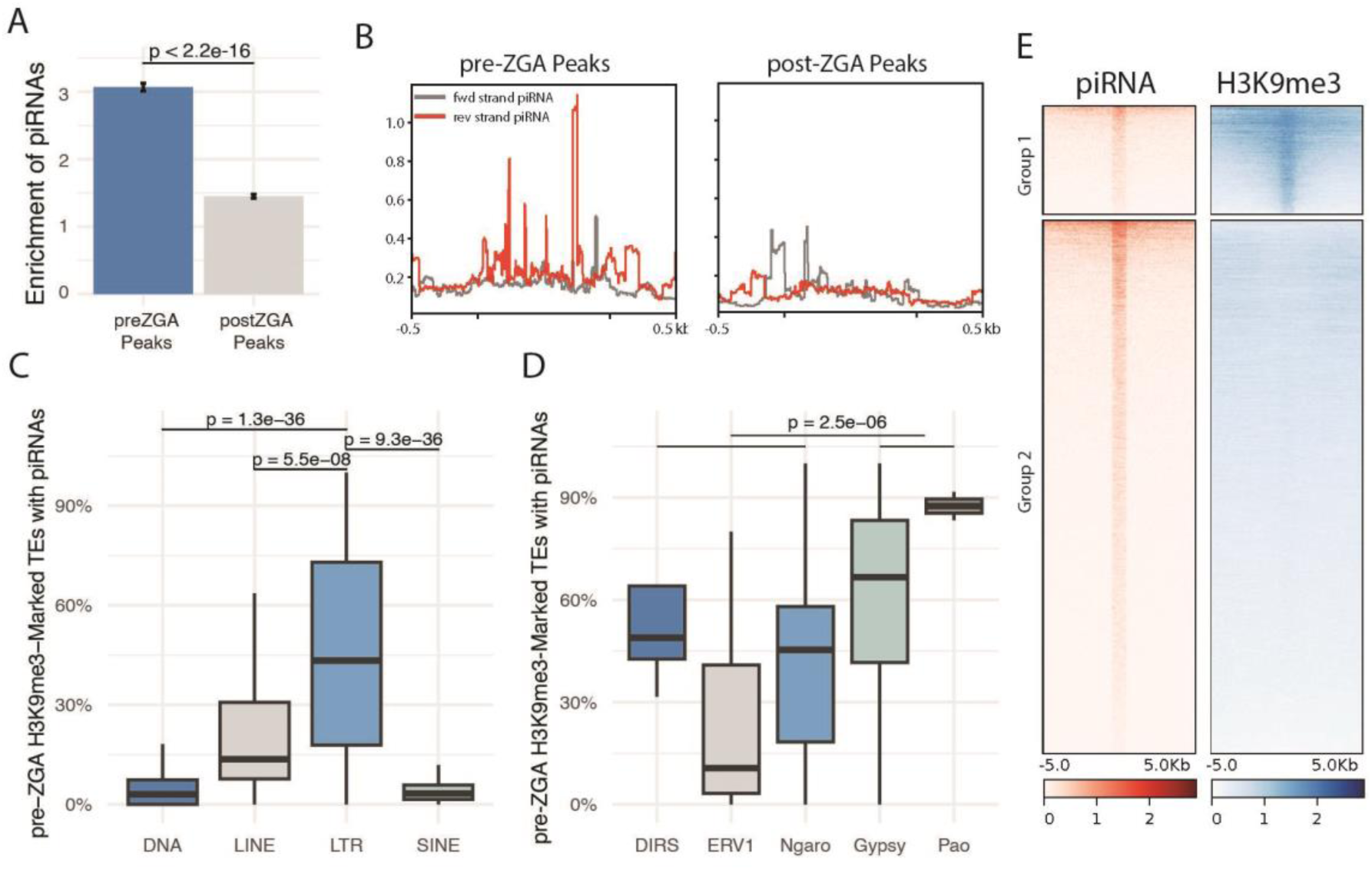
pre-ZGA H3K9me3 is biased towards LTR transposons with enrichment for corresponding maternal piRNAs. **A**. Fold enrichment of piRNAs over control regions (1000 reps) for pre-ZGA H3K9me3 peaks and post-ZGA H3K9me3 peaks. Two-sample t-test. **B**. piRNA signal from the forward (grey) and reverse (red) strands panel across pre-ZGA and post-ZGA peaks (size normalized) with 0.5 kb of flanking region. **C**. Percent of pre-ZGA H3K9me3 enriched elements per transposon family that have associated piRNAs. Two sample t-test. **D**. Identical analysis exclusively examining LTR transposons. **E**. Reverse strand piRNA and 3 hpf H3K9me3 signal at regions of shared enrichment (Group 1) or piRNA enrichment only (Group 2). For all box and whisker plots: center line represents the median, hinges represent 25^th^ and 75^th^ percentiles, and whiskers extend an additional 1.5 times their respective interquartile range. Outliers not plotted. For all panels, only elements with a mappability score of at least 0.1, and families with at least 10 elements meeting this threshold are considered.

### Pericentromeric regions are preferred sites of pre-ZGA H3K9me3

To identify additional characteristics that could delineate pre-ZGA and post-ZGA H3K9me3 enrichment peaks, we next asked whether H3K9me3 peaks that were detected before ZGA were preferentially enriched for specific genomic regions. We noted that many internal regions of high pre-ZGA H3K9me3 peak density overlapped with regions identified as pericentromeric by their BRSATI content (Figure 6A, **S8**). In contrast, post-ZGA H3K9me3 establishment peaks appeared more equally distributed across chromosomes (Figure 6A, **S8**). To quantify this effect, we calculated the fold enrichment of pre-ZGA or post-ZGA peaks in pericentromeric regions compared to 1000 sets of random, size-controlled, non-overlapping null regions from each corresponding chromosome. This analysis confirmed a significantly higher relative chromosomal enrichment of H3K9me3 at pericentromeric regions compared to non-pericentromeric sequences at pre-ZGA timepoints when compared to post-ZGA time points (Figure 6B). Within pericentromeric regions, pre-ZGA peaks overlapped primarily with LTR and DNA transposons, with additional more minor contributions from LINE and SINE elements (Figure 6C).

**Figure 6.**
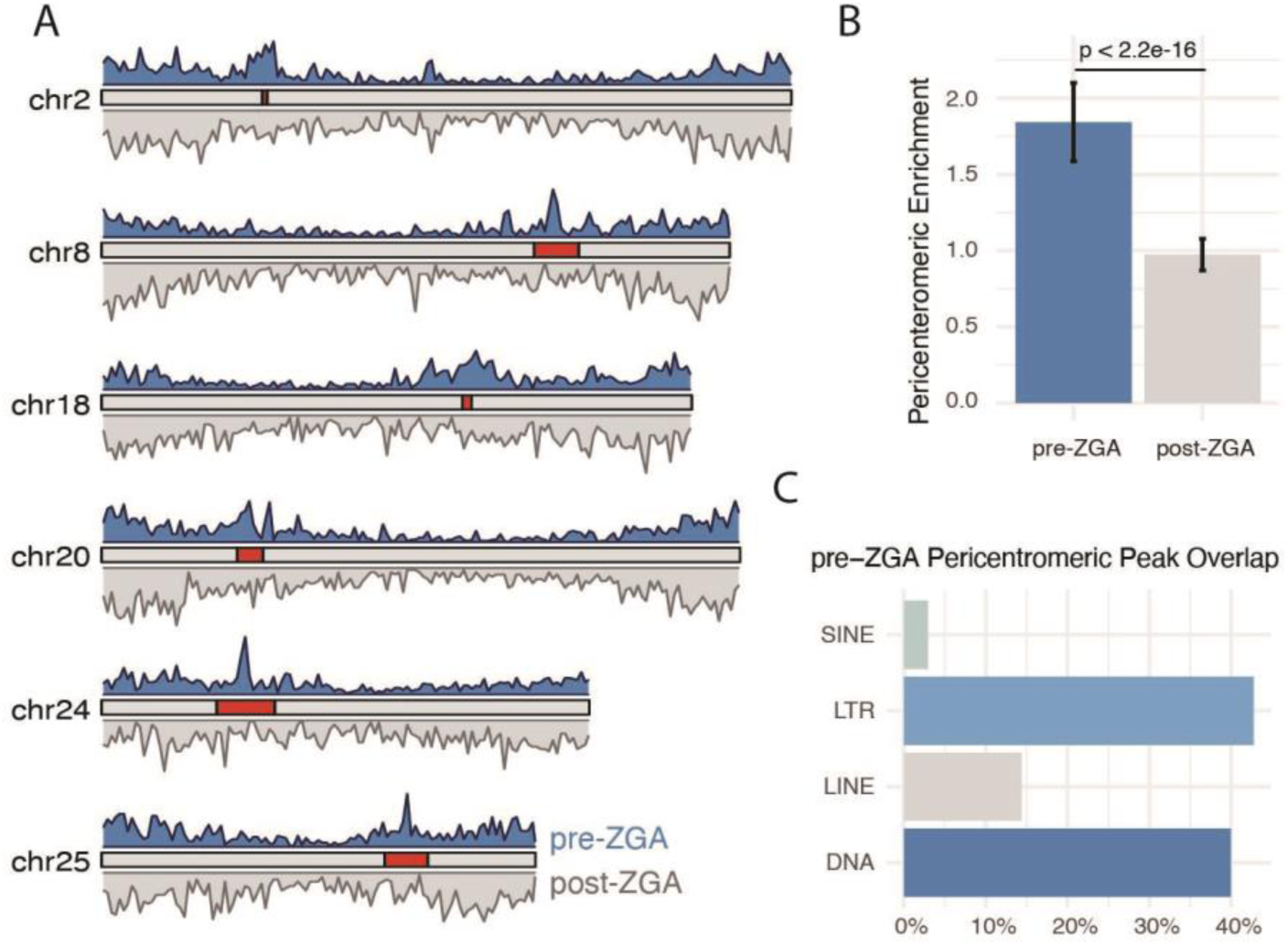
Pre-ZGA peaks are enriched at pericentromeres **A**. Karyoplot showing peak density of pre-ZGA peaks (top, window size = 35 kb) and post-ZGA peaks (bottom, window size = 35 kb) and their location relative to pericentromeric regions (red). **B**. Fold enrichment of pericentromeric intersection over random control regions on each chromosome (1000 reps) for pre-ZGA or post-ZGA H3K9me3 peaks. Two-sample t-test. **C**. Proportion of pre-ZGA pericentromeric peaks overlapping each category of repetitive sequences, only elements with a mappability score of at least 0.1, and families with at least 10 elements meeting this threshold are considered.

### Synergy between piRNA enrichment and pericentromeric localization

To determine if piRNA enrichment and pericentromeric localization might act synergistically to promote pre-ZGA H3K9me3 establishment, we next compared H3K9me3 signal strength at pre-ZGA H3K9me3 peaks that were localized to pericentromeres and enriched for maternal piRNAs to peaks with only one or neither feature. We noted that pericentromeric localization in the absence of corresponding maternal piRNAs did not significantly bias sequences toward stronger pre-ZGA enrichment, suggesting chromosomal location alone is not sufficient to disproportionately promote pre-ZGA H3K9me3. In contrast, among piRNA enriched sequences, those found at pericentromeres showed increased pre-ZGA H3K9me3 enrichment compared to those in other regions of the genome (Figure 7A). Moreover, this synergy persisted, suggesting that early H3K9me3 was predictive of later enrichment (Figure 7B). Similarly, among piRNA associated regions, we noted those in pericentromeres were significantly more likely to exhibit pre-ZGA H3K9me3 peaks compared to those in non-pericentromeric regions (Figure 7C).

**Figure 7.**
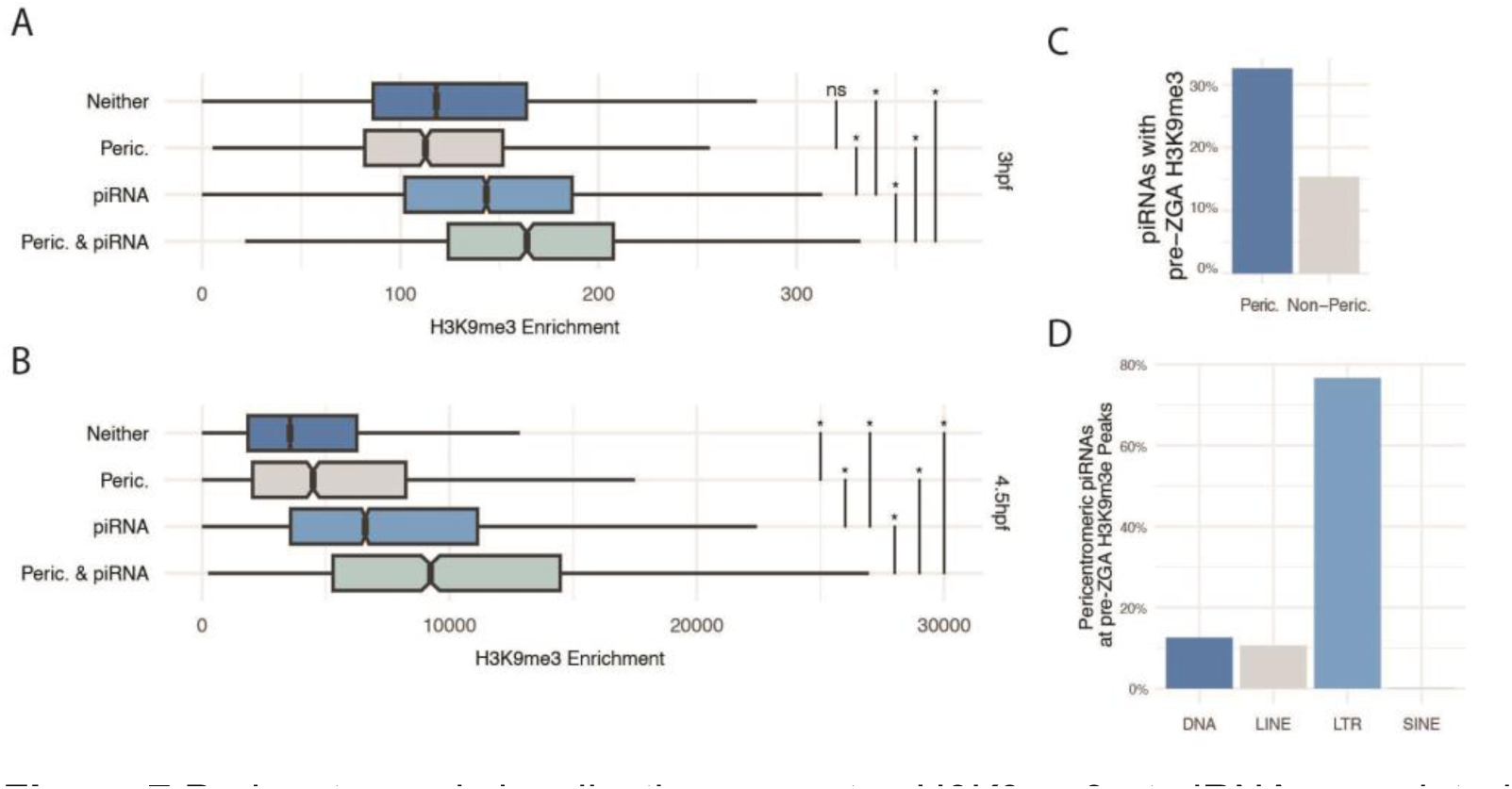
Pericentromeric localization promotes H3K9me3 at piRNA associated sequences. **A**. Average, peak size-normalized, H3K9me3 enrichment at 3 hpf for peaks that overlap with the pericentromere only, piRNAs only, both features, or neither feature. Two sample t-tests, * indicates p <0.05. **B**. Average, peak size-normalized, H3K9me3 signal at 4.5 hpf for peaks established pre-ZGA that overlap with the pericentromere, piRNAs, both, or neither. Two-sample t-tests, * indicates p <0.05. **C**. Fraction of expressed (TPM > 0.5) piRNAs that result in pre-ZGA H3K9me3 enrichment) in either the pericentromere or non-pericentromeric portions of the genome. **D**. Fraction of pre-EGA H3K9me3 peaks associated with piRNA at each transposon category. For all box and whisker plots, center line represents the median, hinges represent 25^th^ and 75^th^ percentiles, and whiskers extend an additional 1.5 times their respective interquartile range. Outliers not plotted.

Not surprisingly, the bulk of sites where synergy was observed corresponded to LTR transposons (Figure 7D). As some LTR sequences are preferentially enriched at pericentromeres, we considered the possibility that the impact of pericentromeres could be explained simply as function of transposon localization. However, direct comparison of pre-ZGA H3K9me3 at LTR elements within vs outside pericentromeric regions, again revealed significant bias toward pericentromeres, particularly for Gypsy and Pao elements (**Figure S9**). Together, these findings suggest that pericentromeric regions may provide a more favorable environment for successful early targeting by the piRNA machinery.

### H3K9me3 patterns continue to evolve during later embryogenesis

Finally, we investigated how well H3K9me3 patterns laid down in the pluripotent blastula correspond to those at later stages of embryonic development. For this analysis, we compared the H3K9me3 landscape of 4.5 hpf embryos to that of 24 hpf embryos using CUT&RUN. We noted substantial maturation of the heterochromatic landscape over this period. At 24 hpf, we found that across the genome, H3K9me3 remained predominantly enriched at repetitive sequences, but the average signal strength was reduced, and the specific repertoire of enriched transposons shifted (Figure 8A-B, **S10**). Between 4.5 hpf and 24 hpf embryos, H3K9me3 enrichment was shared at ∼41,000 sites, while another ∼30,000 sites were unique to each time point (Figure 8A-B). H3K9me3 sites gained in 24 hpf embryos were biased toward DNA transposons, including hAT-Ac, PIF-Harbinger, and DNA families, suggesting that these transposons may be targeted for silencing later in development by late embryo specific or cell -type specific pathways (Figure 8C, **E**). In contrast, LTRs, and specifically Gypsy and DIRS elements, are overrepresented among elements that lose H3K9me3 enrichment by 24 hpf (Figure 8C-D). This observation is consistent with the idea that cell type or developmental stage specific mechanisms help drive an evolving H3K9me3 landscape.

**Figure 8.**
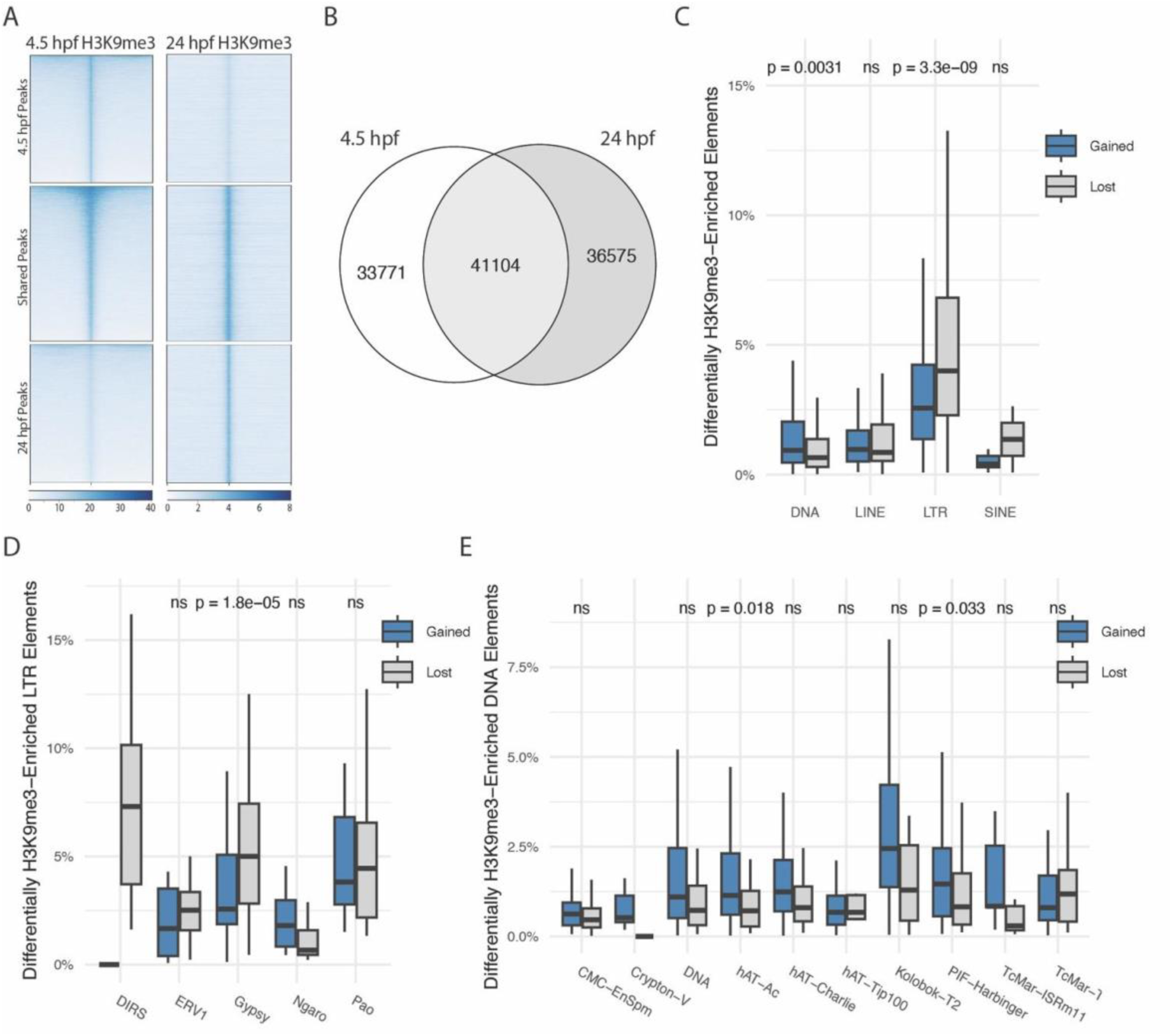
The H3K9me3 landscape shifts across late embryogenesis. **A**. Heat map depicting regions with H3K9me3 enrichment specific to 4.5 hpf, 24 hpf, or both time points. **B**. Venn diagram representing the intersection of H3K9me3 peaks between 4.5 hpf and 24 hpf. **C**. Percent of each element type that gains or loses H3K9me3 between 4.5 hpf and 24 hpf. Two-sample t-test. **D**. Percent of each LTR element type that gains or loses H3K9me3 enrichment between 4.5 hpf and 24 hpf. Two-sample t-test. No DIRS families gained H3K9me3. **E**. Percent of each DNA element type that gains or loses H3K9me3 enrichment between 4.5 hpf and 24 hpf. Two-sample t-test. No Cypton-V families lost H3K9me3. For all box and whisker plots: center line represents the median, hinges represent 25^th^ and 75^th^ percentiles, and whiskers extend an additional 1.5 times their respective interquartile range. Outliers not plotted. For all panels, only elements with a mappability score of at least 0.1, and families with at least 10 elements meeting this threshold are considered.

## Discussion

Collectively, our results provide a comprehensive assessment of the emerging H3K9me3 landscape across zebrafish embryogenesis, uncovering distinct features of pre- and post-ZGA H3K9me3 establishment, as well as further maturation of H3K9me3 enrichment profiles during later embryonic development. We identify sequences corresponding to maternal piRNAs with pericentromeric localization as favored sites of pre-ZGA H3K9me3 enrichment, and we uncover a striking jump in the amplitude of pre ZGA H3K9me3 peaks, over just 30 min corresponding to the onset of ZGA.

Our detection of low-amplitude pre-ZGA H3K9me3 enrichment in the zebrafish embryo builds on publications noting low levels of pre-ZGA H3K9me3 establishment in the mouse paternal pronucleus by immunofluorescence, and H3K9me3 at a small subset of sequences in stage 3 Drosophila embryos by ultra-low input ChIP (Burton et al. 2020; Wei et al. 2021). Our results combine with these studies to suggest pre-ZGA H3K9me3 establishment at some sequences may be a conserved feature in animal embryogenesis.

While time course analysis revealed linear growth in the number of H3K9me3 peaks detected over time, quantitative comparison of signal amplitude across time points instead showed a dramatic surge in H3K9me3 over just 30 minutes coinciding with the onset of the major wave of embryonic transcription.

Increased peak amplitudes are most easily explained by an increase in the fraction of cells harboring H3K9me3 at a given site, which is consistent with a model in which early establishment involves stochasticity. Such stochasticity could arise from a need for repeated *de novo* targeting of H3K9me3 during every round of cell division without inheritance, or from competition between mechanisms promoting H3K9me3 establishment and antagonistic mechanisms blocking establishment or promoting H3K9me3 removal (Laue et al. 2019; Burton et al. 2020). There is also potential that the abbreviated length of cell divisions in the early zebrafish embryo impacts H3K9me3 levels by limiting the window of time available for establishment. This latter model is supported by observations that reducing cell cycle length impeded marks of heterochromatin formation in Drosophila embryos (Seller et al. 2019).

Among H3K9me3 peaks established before ZGA, we observed a strong preference for LTR transposons, with Gypsy and Pao elements showing the strongest bias toward pre-ZGA enrichment. Similar biases were not observed for tyrosine recombinase encoding DIRS and Ngaro elements, suggesting distinct mechanisms may be involved in the targeting of these elements, which are thought to be mostly absent from Drosophila and mammalian genomes. Intriguingly, among the 8 families of DNA elements exhibiting substantial H3K9me3 enrichment, we note three are tyrosine recombinase encoding Cryptons, raising the possibility that tyrosine recombinase sequences themselves may help identify these elements as foreign.

Mechanistically, our data suggest that much of pre-ZGA targeting of zebrafish LTR elements is driven by maternally loaded piRNAs. This finding builds on observations reporting the enrichment of embryonic piRNAs in genomic regions near some sites of early H3K9me3 establishment in Drosophila and the detection of maternally loaded piRNAs corresponding to LTR transposons that are repressed by H3K9me3 in early gastrula stage embryos in zebrafish (Guo et al. 2021; Wei et al. 2021). Together, these findings suggest that piRNAs may have generalized roles outside the germline in regulating early H3K9me3 establishment at some transposon sequences. Alongside piRNAs, we also identified pericentromeric localization as a strong predictor of pre-ZGA H3K9me3 establishment, with piRNA associated sequences within pericentromeres significantly more likely to exhibit pre-ZGA H3K9me3 enrichment compared to those outside of these regions. It is possible that the increased density of repetitive sequences within these regions provides some level of stabilization to early stochastic targeting events.

It is also likely that transcription-dependent mechanisms of H3K9me3 regulation further contribute to the shifting patterns of H3K9me3 that we observe following ZGA. For example, although zebrafish lack KRAB-ZF proteins, an important component of H3K9me3 targeting in tetrapod animals, they do encode for an alternative group of zinc finger proteins called FiNZ-ZNF proteins that may play a similar role in zebrafish and are first expressed at ZGA (Rosspopoff and Trono 2023; Wells et al. 2023). It is also possible that post-ZGA transcription of transposons themselves may generate RNAs that are recognized by host machinery to drive their silencing (Almeida et al. 2022). We anticipate that the high-resolution, low background time course data generated in this study will provide a strong foundation for defining additional inputs that promote directed H3K9me3 silencing across early development.

## Data Availability

All sequencing data generated in this study including initial H3K9me3 CUT&RUN time course and the follow up CUT&RUN experiments at 2.5 hpf, 4.5 hpf, and 24 hpf with Abcam, Diagenode, and Active Motif H3K9me3 antibodies, is available for download from GEO (GSE256288) and a detailed overview of sequencing information is available in **Table S1**. Code used for data analysis and figure generation is publicly available at https://github.com/Goll-Lab/H3K9me3_Kinetics_2024.

## Supporting information

manuscript file

## Acknowledgements

We thank Garren Davis, Tod Butenschon, and University Research Animal Resources at UGA for zebrafish husbandry and support. This study was supported in part by resources and technical expertise from the Georgia Advanced Computing Resource Center and the Georgia Genomics and Bioinformatics Core.

