## Supplementary material for "The emerging H3K9me3 chromatin landscape during zebrafish embryogenesis": manuscript file

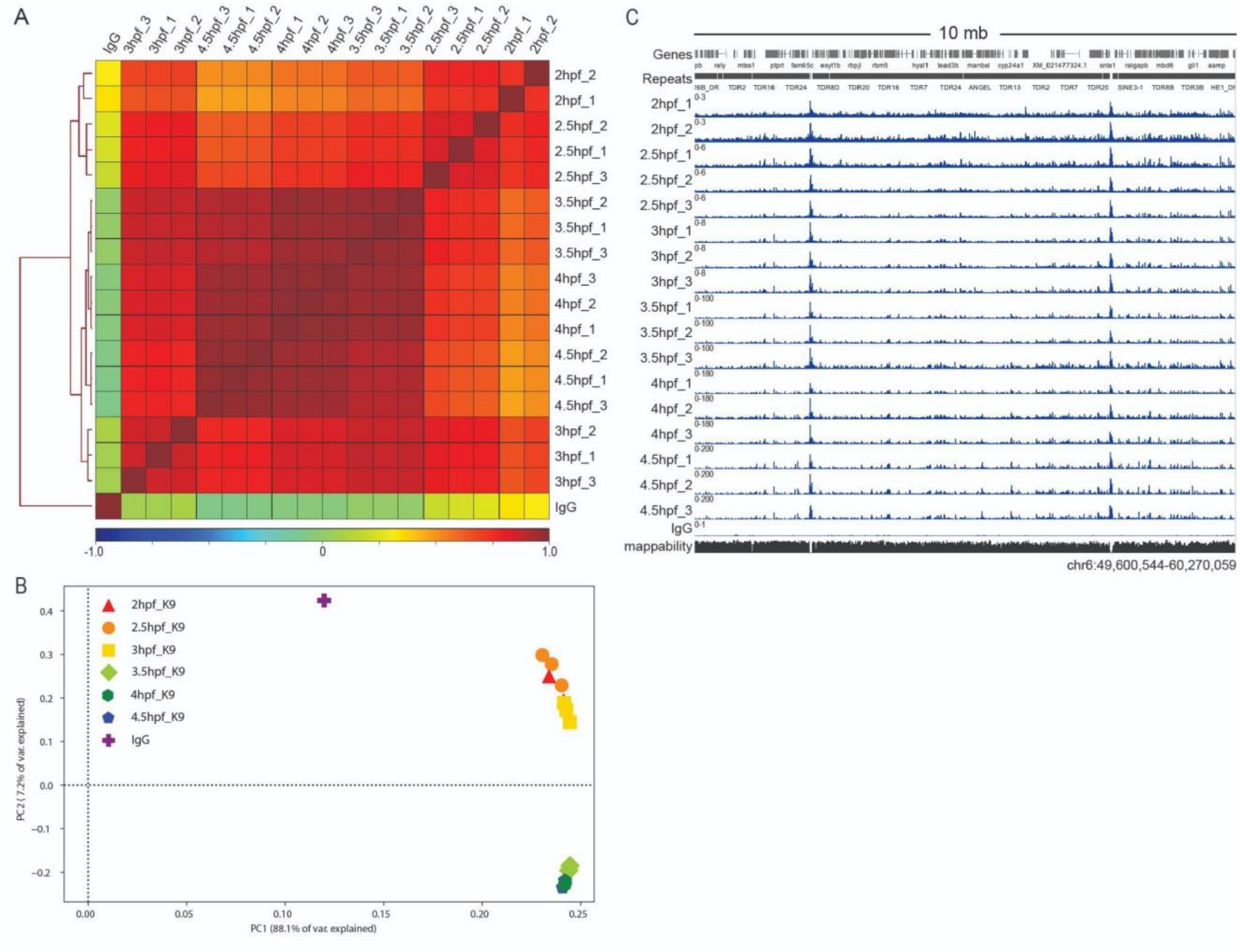

**Figure S1**. High correlation across CUT&RUN sample replicates at all profiled developmental stages. **A**. Spearman correlation heatmap demonstrating high correlation between CUT&RUN replicates at each time point. IgG is averaged across replicates from all timepoints. **B**. PCA analysis of CUT&RUN replicates. IgG is averaged across replicates from all timepoints. **C**. Genome browser image demonstrating strong concordance in signal between CUT&RUN sample replicates.

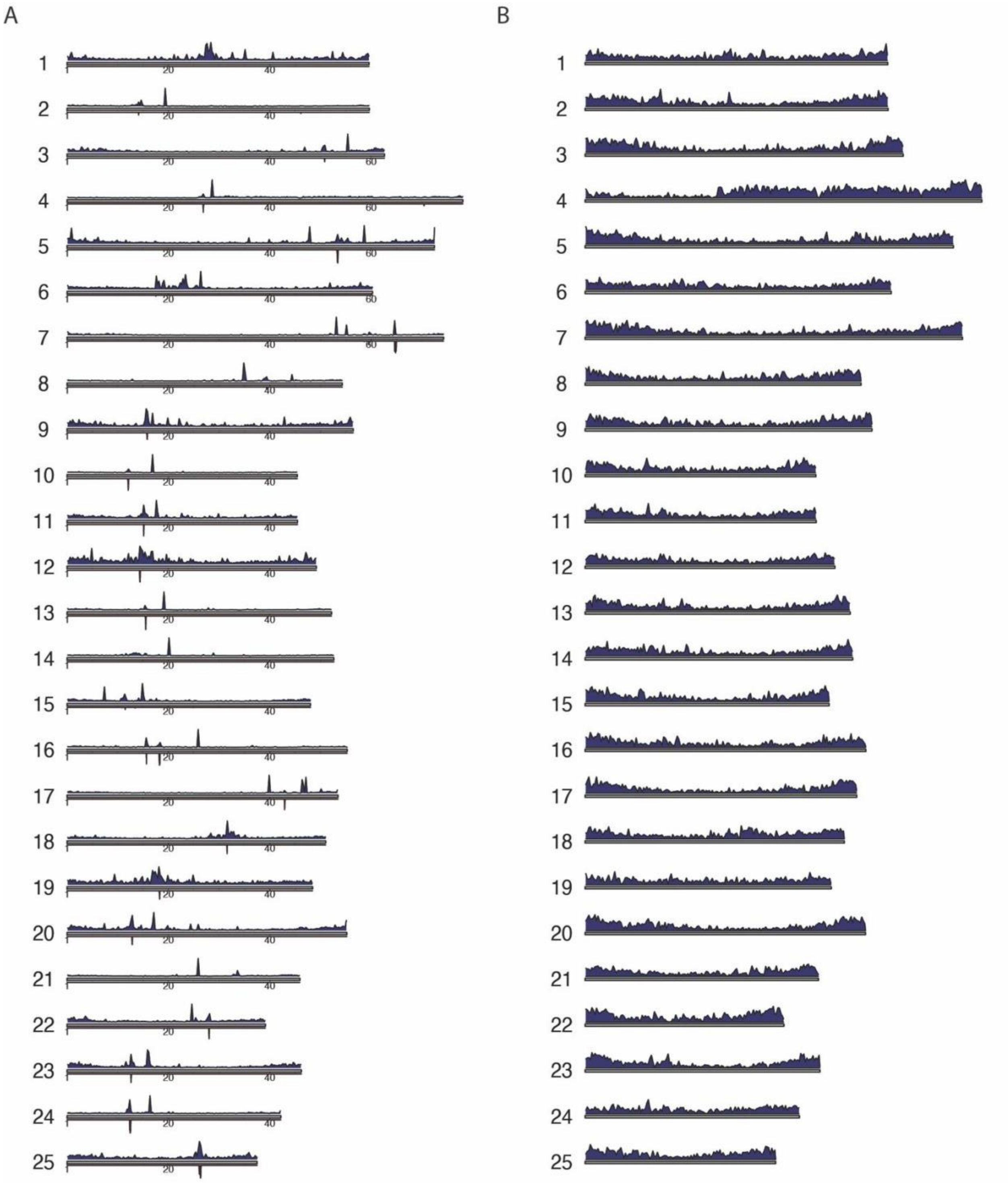

**Figure S2**. Genome-wide chromosomal landscape of H3K9me3 at 4.5 hpf **A**. Karyoplot of average 4.5 hpf H3K9me3 coverage (top, bin size = 250 kb) and pericentromeric repeat BRSATI density (bottom, window size = 50 kb) across all 25 zebrafish chromosomes. **B**. Karyoplot of 4.5 hpf H3K9me3 peak density (window size = 30 kb) across all 25 zebrafish chromosomes.

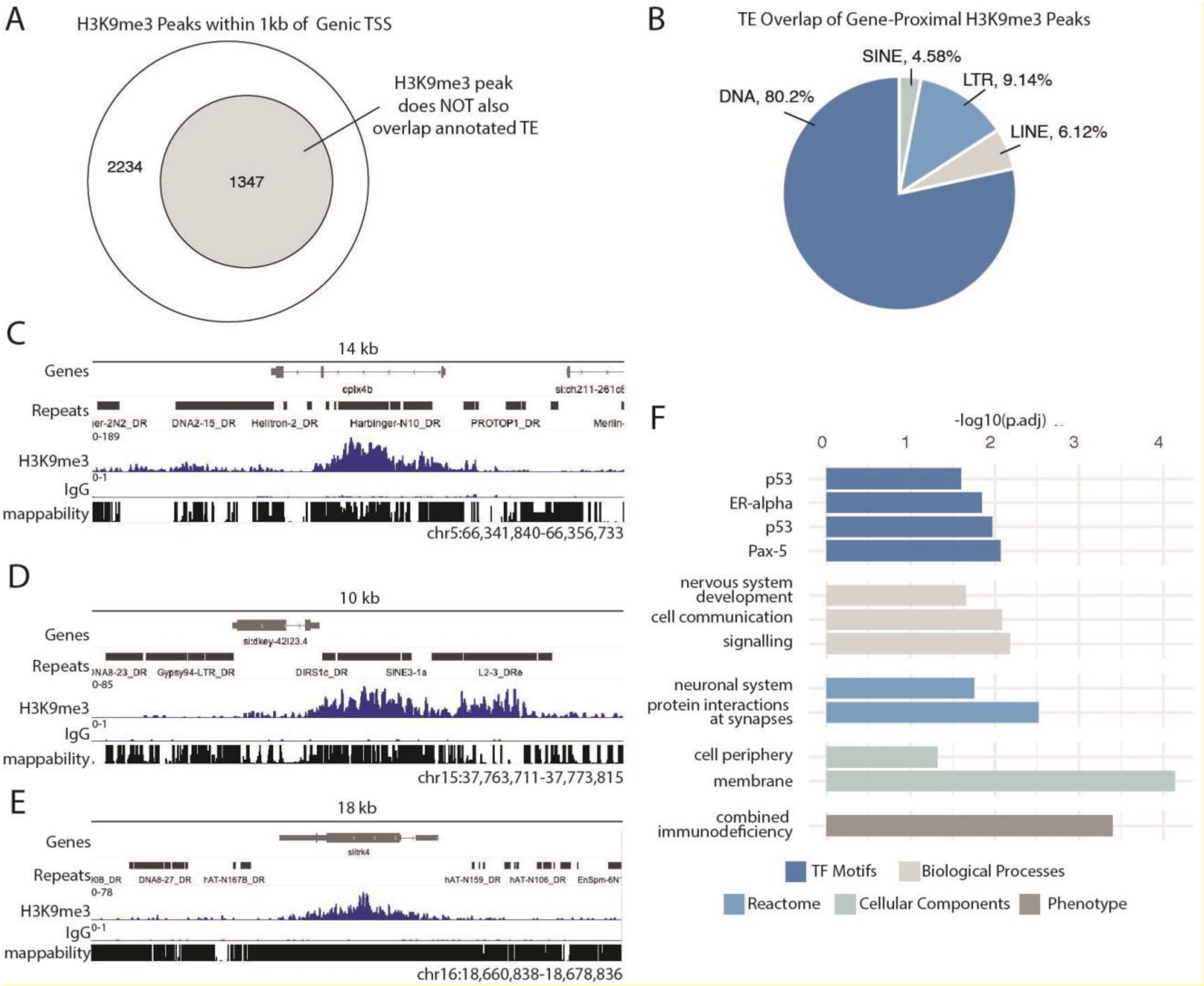

**Figure S3**. Genic H3K9me3. **A**. Categorization of H3K9me3 peaks within 1kb of a genic TSS. **B**. Fraction of gene-proximal transposable elements enriched for H3K9me3 for each transposon category. **C-E**. Genome browser image showing an example of 4.5 hpf H3K9me3 enrichment within a gene but also overlapping an intronic TE (**C**), directly upstream of genic TSS but also overlapping a TE (**D**) and within a gene and not overlapping a TE (**E**). **F.** Significant terms from gene ontology analysis of protein coding genes with H3K9me3 enrichment with 1 kb of TSS. For all panels, data represents 4.5 hpf H3K9me3 signal averaged across replicates and IgG signal averaged replicates from all timepoints.

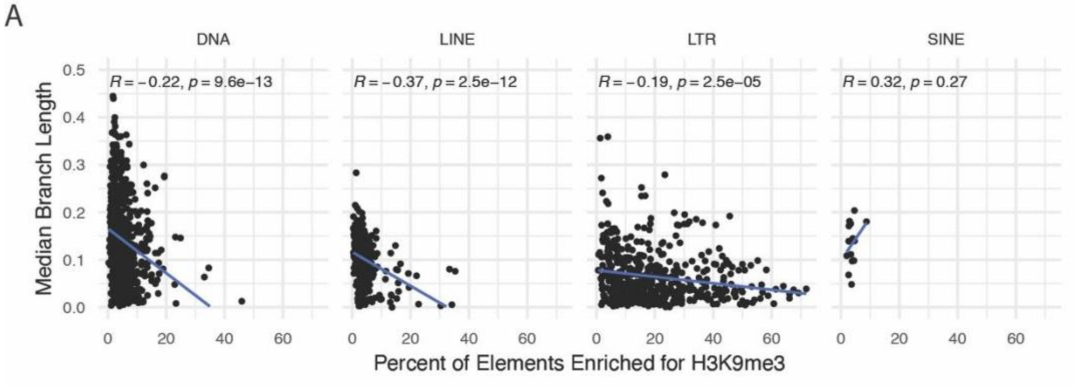

**Figure S4**. H3K9me3 enrichment correlated with transposon age. **A.** Percent elements of each transposon family enriched for H3K9me3 correlates with transposon median phylogenetic branch length as calculated by Chang et al 2022. Pearson correlation calculation for coefficient and p value.

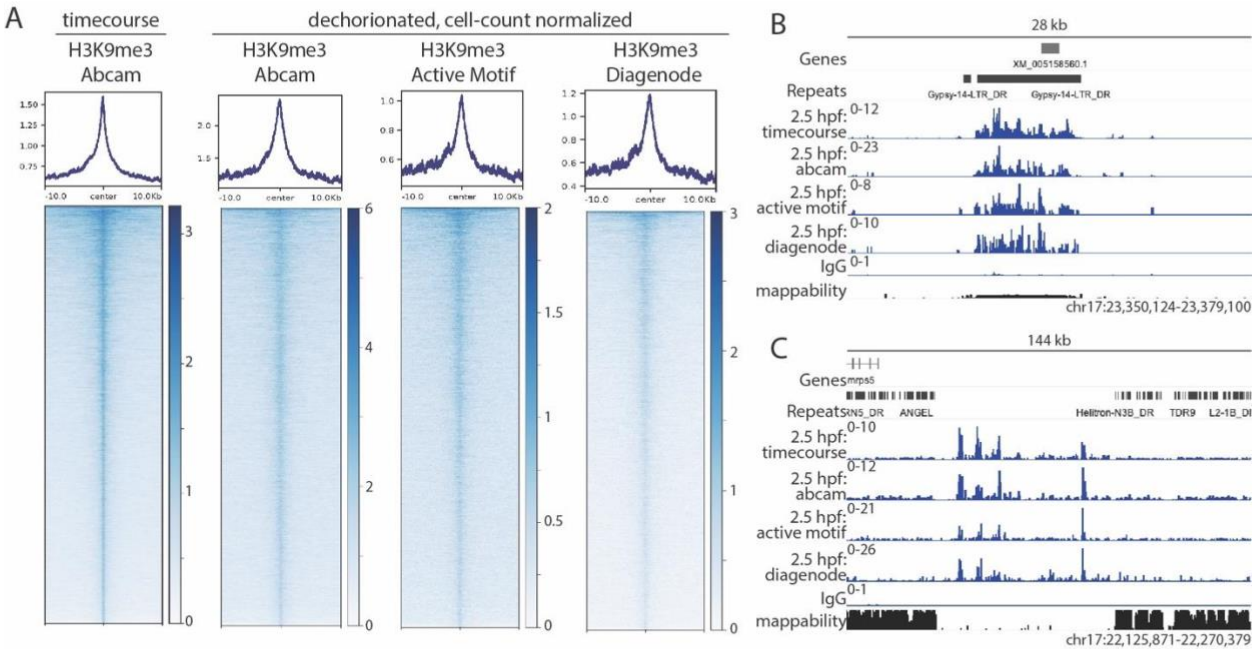

**Figure S5**. 2.5 hpf H3K9me3 enrichment detectable by multiple antibodies in dechorionated samples. **A**. 2.5 hpf H3K9me3 peaks from original time course are also enriched for H3K9me3 in dechorionated, cell-count normalized samples with three commercially produced antibodies. Regions in heatmap are all ordered the same, based on signal intensity in the Abcam timecourse. **B** & **C**. Genome browser images showing high correlation between H3K9me3 signal in all four 2.5 hpf H3K9me3 CUT&RUN experiments. For all panels, data represents H3K9me3 signal averaged across replicates and IgG signal averaged across replicates.

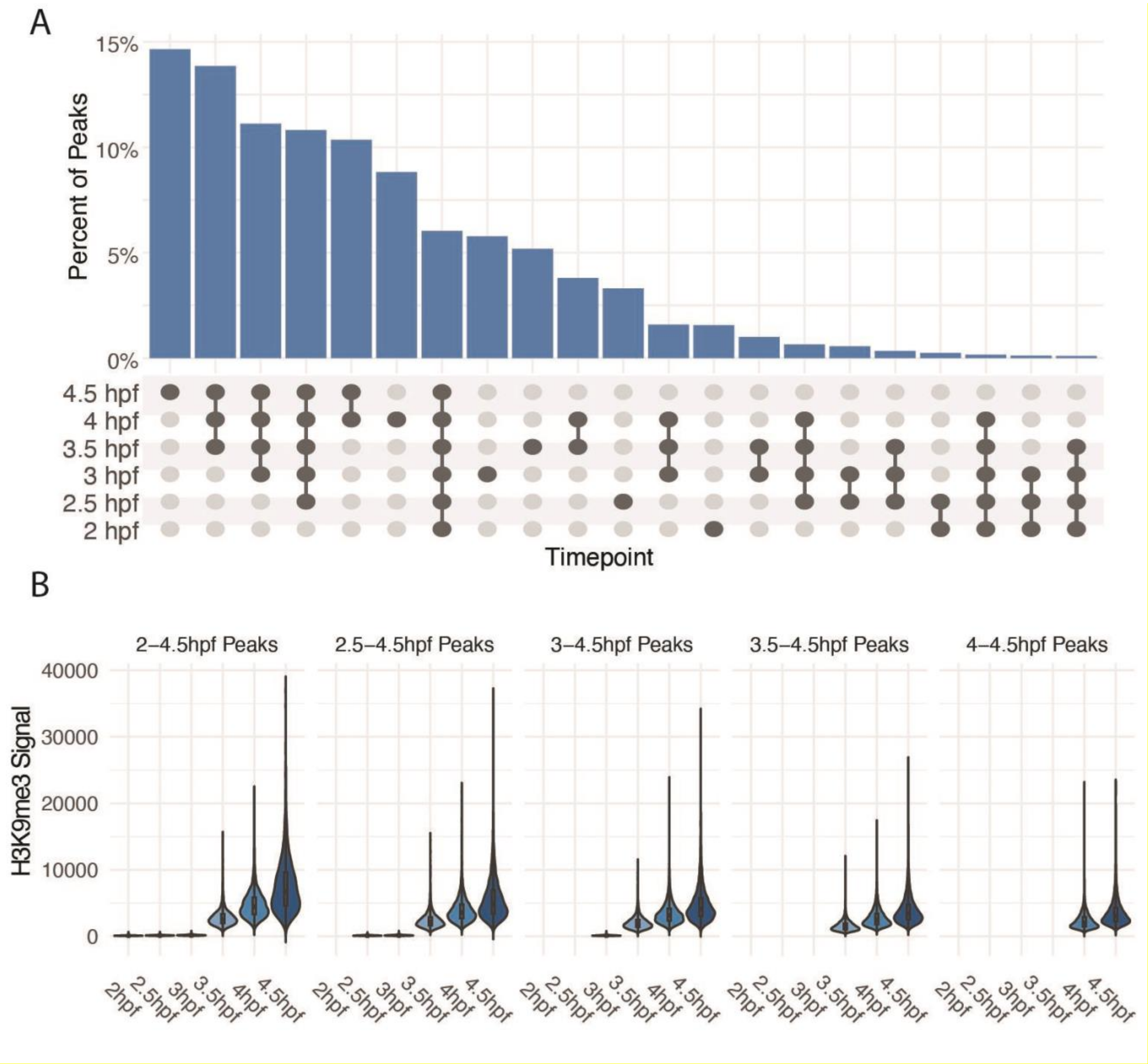

**Figure S6**. H3K9me3 peaks detected in the early embryo persist across development. **A**. Upset plot demonstrating H3K9me3 peak overlap across all six timepoints in the CUT&RUN developmental time course. Peaks that fell into multiple non-consecutive timepoints not included. **B**. Average H3K9me3 signal for peaks that initiate at each timepoint and persist through 4.5 hpf, values outside 5 standard deviations not plotted for ease of visualization.

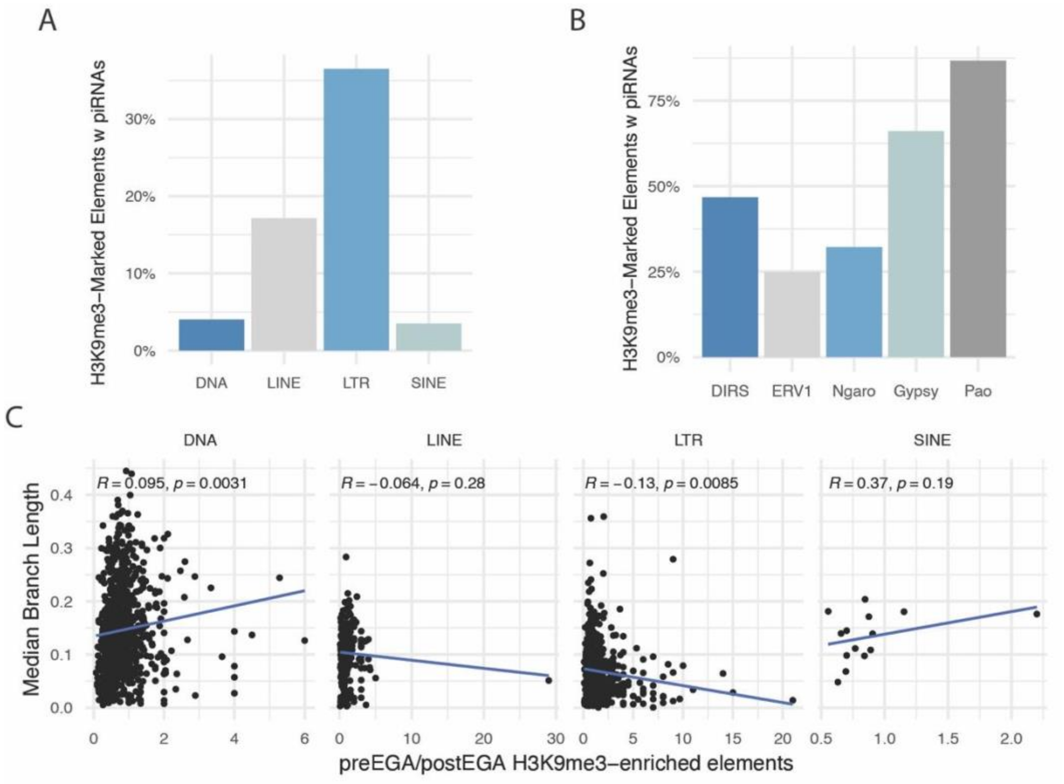

**Figure S7**. pre-ZGA H3K9me3 enrichment **A**. Percent of pre-ZGA H3K9me3 enriched elements that have associated piRNAs. **B**. Percent of pre-ZGA H3K9me3 enriched elements that have associated piRNAs for LTR families. **C**. Ratio of H3K9me3-enriched pre-ZGA elements to H3K9me3-enriched post-ZGA elements for each transposon family correlates with median phylogenetic branch length as calculated by Chang et al 2022. Pearson correlation calculation for coefficient and p value.

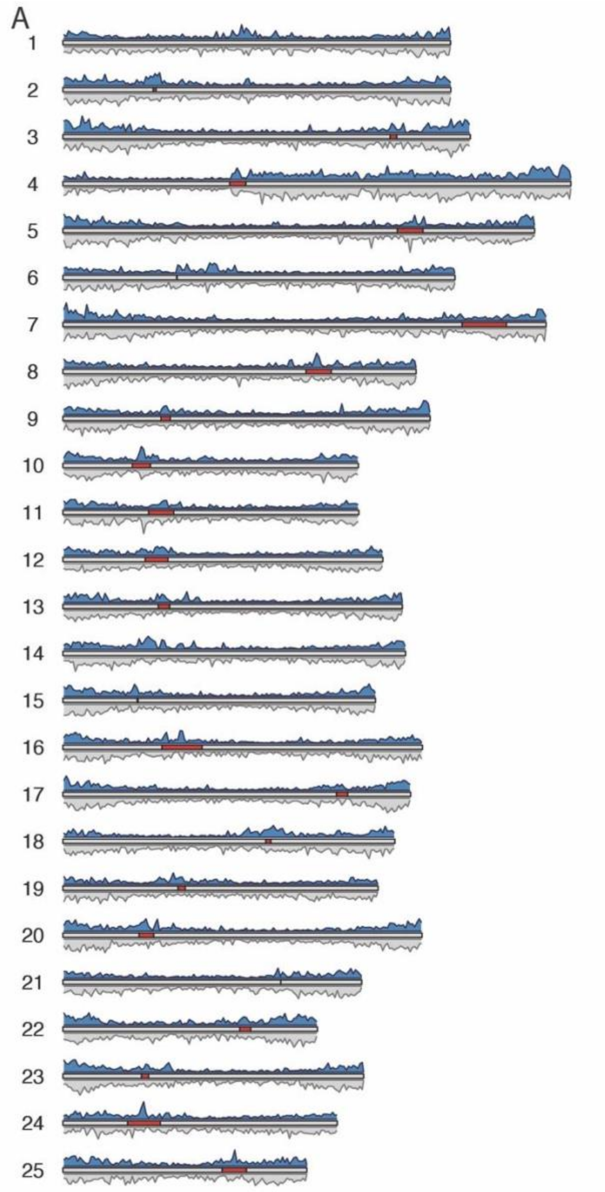

**Figure S8**. pre-ZGA H3K9me3 peaks enriched near pericentromeres. **A**. Peak density of pre-ZGA peaks (top, window size = 35 kb) and post-ZGA peaks (bottom, window size = 35 kb) across zebrafish. Pericentromeric regions denoted in red.

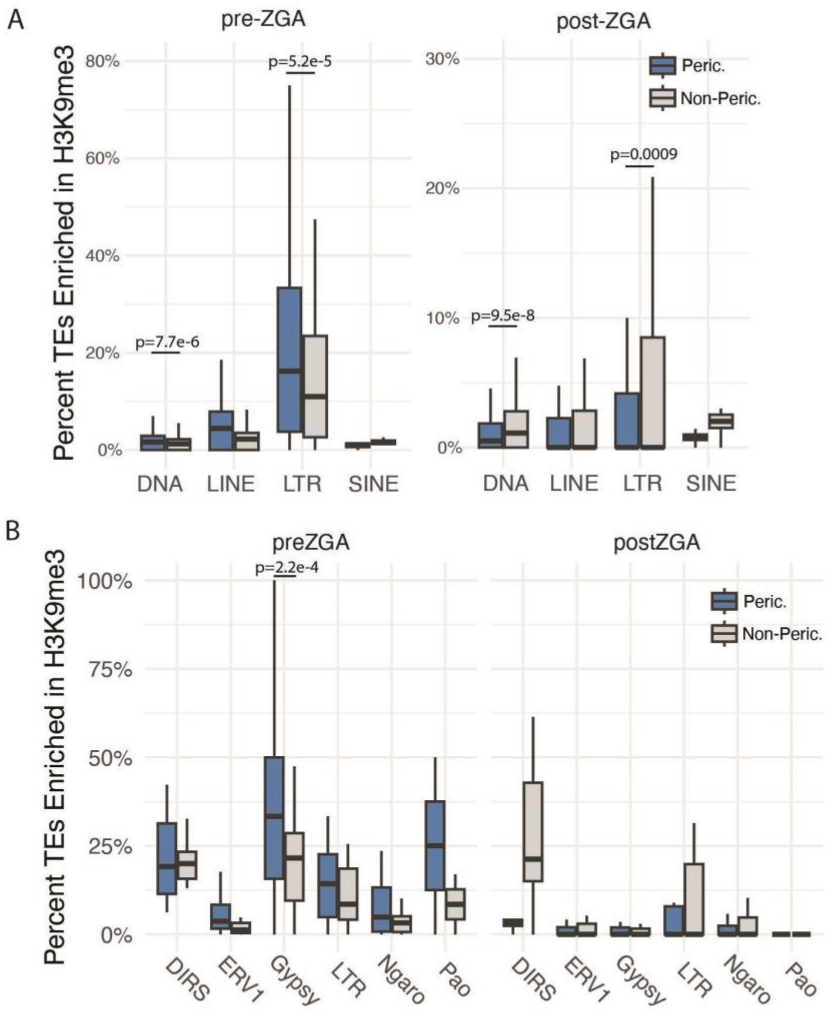

**Figure S9. A**. Percent of pericentromeric or non-pericentromeric elements with H3K9me3 established pre- or post-ZGA per transposon family for each category. Two sample t-test. **B**. Percent of pericentromeric or non-pericentromeric elements with H3K9me3 established pre- or post-ZGA per transposon family for each LTR category. Two sample t-test. For all box and whisker plots, center line represents the median, hinges represent 25$^{th}$ and 75$^{th}$ percentiles, and whiskers extend an additional 1.5 times their respective interquartile range. Outliers not plotted.

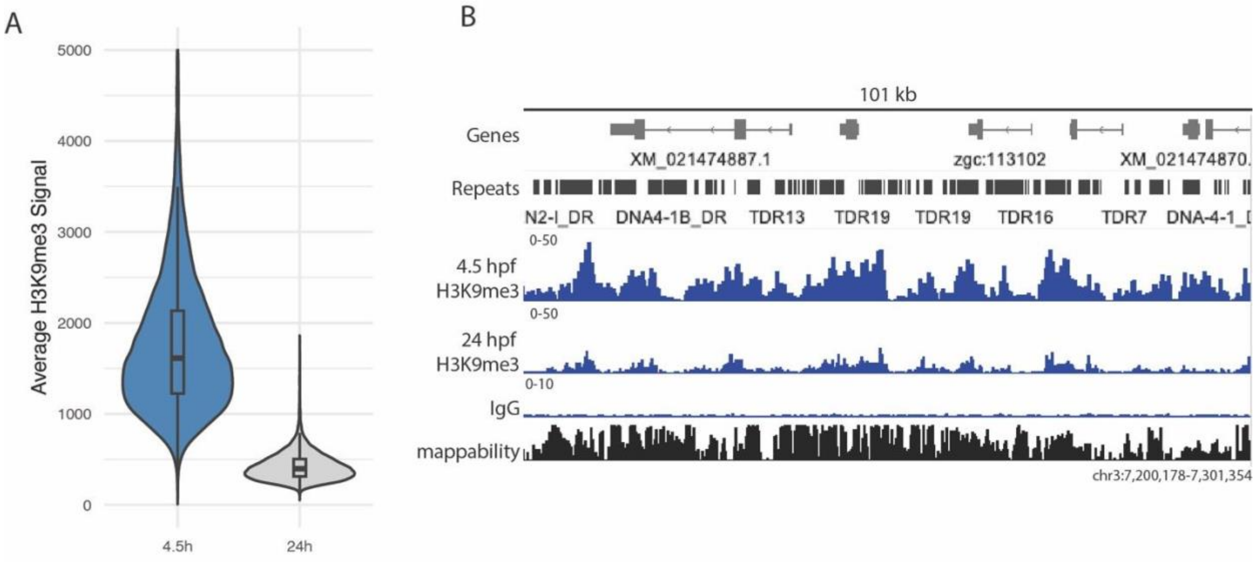

**Figure S10**. H3K9me3 signal reduction from 4.5 hpf to 24 hpf. **A**. Average H3K9me3 signal in peaks shared between 4.5 hpf and 24 hpf at each timepoint. **B**. Genome browser image example of a region with H3K9me3 signal depletion at 24 hpf compared to 4.5 hpf. For all panels, data represents H3K9me3 signal averaged across 3 replicates for each timepoint and IgG signal averaged across replicates from all timepoints.

| Sample | Trimmed Reads | Aligned Reads | Percent Aligned |
|---|---|---|---|
| *Time Course* | | | |
| 2hpf_IgG_1 | 7084738 | 2356995 | 33.27% |
| 2hpf_K9_1 | 9784762 | 5095451 | 52.08% |
| 2hpf_K9_2 | 10755816 | 5879744 | 54.67% |
| 2.5hpf_IgG_1 | 6031753 | 2003168 | 33.21% |
| 2.5hpf_ IgG _2 | 6305478 | 2051922 | 32.54% |
| 2.5hpf_K9_1 | 11774745 | 6606392 | 56.11% |
| 2.5hpf_K9_2 | 15844530 | 8997902 | 56.79% |
| 2.5hpf_K9_3 | 12952050 | 6980894 | 53.90% |
| 3hpf_ IgG _1 | 3866283 | 1310690 | 33.90% |
| 3hpf_ IgG _2 | 5142553 | 1433110 | 27.87% |
| 3hpf_K9_1 | 10234396 | 5410725 | 52.87% |
| 3hpf_K9_2 | 9913319 | 5281533 | 53.28% |
| 3hpf_K9_3 | 11282709 | 4494363 | 39.83% |
| 3.5hpf_ IgG _1 | 6517397 | 3428167 | 52.60% |
| 3.5hpf_K9_1 | 24473518 | 13343339 | 54.52% |
| 3.5hpf_K9_2 | 31516460 | 17126515 | 54.34% |
| 3.5hpf_K9_3 | 23561722 | 12886808 | 54.69% |
| 4hpf_ IgG _1 | 6783381 | 3581281 | 52.79% |
| 4hpf_ IgG _2 | 7267968 | 3698412 | 50.89% |
| 4hpf_K9_1 | 44389054 | 23226239 | 52.32% |
| 4hpf_K9_2 | 34135770 | 18056114 | 52.89% |
| 4hpf_K9_3 | 32187842 | 16992055 | 52.79% |
| 4.5hpf_ IgG _1 | 10742276 | 5522664 | 51.41% |
| 4.5hpf_ IgG _2 | 9726450 | 5261038 | 54.09% |
| 4.5hpf_K9_1 | 37342412 | 18288186 | 48.97% |
| 4.5hpf_K9_2 | 29126156 | 14243527 | 48.90% |
| 4.5hpf_K9_3 | 31550834 | 15331219 | 48.59% |
| *2.5 hpf Confirmation* | | | |
| K9abcam_2.5hpf_1 | 9456915 | 5734770 | 60.64% |
| K9abcam_2.5hpf_2 | 10001717 | 5845900 | 58.45% |
| K9abcam_2.5hpf_3 | 4687171 | 2699017 | 57.58% |
| K9active_2.5hpf_1 | 5335975 | 2221699 | 41.64% |
| K9active_2.5hpf_2 | 5556380 | 2252676 | 40.54% |
| K9active_2.5hpf_3 | 3989203 | 1386662 | 34.76% |
| K9dia_2.5hpf_1 | 5194396 | 1695748 | 32.65% |
| K9dia_2.5hpf_2 | 5175525 | 1444652 | 27.91% |
| K9dia_2.5hpf_3 | 2597248 | 728999 | 28.07% |
| IgG_2.5h | 8471424 | 1591798 | 18.79% |
| *24 hpf Analysis* | | | |
| K9abcam_4.5hpf_1 | 8882100 | 5260550 | 59.23% |
| K9abcam_4.5hpf_2 | 12897319 | 7584072 | 58.80% |
| K9abcam_4.5hpf_3 | 11888327 | 7004930 | 58.92% |
| K9abcam_24hpf_1 | 15146177 | 8623185 | 56.93% |
| K9abcam_24hpf_2 | 8026668 | 4523763 | 56.36% |
| K9abcam_24hpf_3 | 7810482 | 1791607 | 22.94% |
| IgG_4.5h | 3545678 | 862640 | 24.33% |
| IgG_24h | 2705651 | 243582 | 9.00% |

**Table S1** Overview of sequencing information used in this study.
